# Charting the undiscovered metabolome with synthetic multiplexing implicates ibuprofen-carnitine in myotoxicity

**DOI:** 10.1101/2025.11.18.689170

**Authors:** Abubaker Patan, Shipei Xing, Vincent Charron-Lamoureux, Zhewen Hu, Victoria Deleray, Julius Agongo, Jasmine Zemlin, Harsha Gouda, Prajit Rajkumar, Yasin El Abiead, Aubreyana E. McMaugh, Hannah Heath, Helena Mannochio-Russo, Ipsita Mohanty, Laila Abolfathi, Ricardo Almada-Monter, Jia Yang, Carlynda Lee, Daniel Leanos, Noah Weimann, Wataru Tsuda, Sadie Giddings, Tammy Bui, Eric Ding, Kine Eide Kvitne, Haoqi Nina Zhao, Simone Zuffa, Paulo Wender Portal Gomes, Vivian Nguyen, Aileen Andrade, Maria A Pawlowski, Ashley C Ferland, Elisabeth Orozco, Wilhan Donizete Gonçalves Nunes, Andrés M. Caraballo-Rodríguez, Lurian Caetano David, Kathleen M. Giacomini, Adrian Jinich, Jeremy Carver, Nuno Bandeira, Mingxun Wang, Lindsey A. Burnett, Dionicio Siegel, Pieter C. Dorrestein

## Abstract

Most molecular features detected in untargeted metabolomics remain uncharacterized due to the limited scope of existing spectral reference libraries. We synthesized >100,000 biologically inspired compounds using multiplexed reactions, of which 91% were absent from existing structural databases, and searched the resulting MS/MS library across >1.7 billion public spectra, increasing annotation rates by 17.4%. This approach revealed previously undescribed exposure-derived metabolites, including ibuprofen–carnitine. Because ibuprofen has been linked to rhabdomyolysis, reduced mitochondrial function, and impaired muscle recovery in carnitine-limited contexts, we investigated the functional relevance of this conjugate. Ibuprofen–carnitine reduced carnitine transport via the OCTN2 transporter, and in a postpartum mouse muscle injury model, ibuprofen delayed muscle repair that could be rescued by carnitine supplementation, with urinary ibuprofen–carnitine:carnitine ratios tracking this effect. These findings support a hypothesis whereby NSAID–carnitine conjugates compete for carnitine transport, impairing energy metabolism and muscle recovery in susceptible individuals. Synthetic multiplexing thus provides a scalable route to annotate the dark metabolome and generate experimentally testable biological hypotheses.

## Main

Tandem mass spectrometry (MS/MS) spectral reference libraries are essential tools in untargeted metabolomics, enabling researchers to propose plausible structural hypotheses for MS/MS of detected ions of metabolites. Although 93.1% of public MS/MS spectra that are not currently annotated include ion forms such as in-source fragments, different adducts, multimers, or chimerics and low information content spectra (e.g., low signal to background or high intensity but few ions containing spectra)^1–3^, the sheer amount of data that remain uncharacterized in publicly deposited metabolomics data likely indicates a significant reservoir of undiscovered biochemistry and uncharacterized metabolic pathways^1,2^. To enable this expansion of metabolite annotation for which standards are available, we constructed an MS/MS spectral reference library through multiplexed organic synthesis. Reactions were conducted on pools of biologically relevant starting materials, and the resulting mixtures were analyzed via liquid chromatography tandem mass spectrometry (LC-MS/MS) to obtain a synthetic reference library where the MS/MS have known structures. This allowed us to assess whether such molecules had been previously observed in public datasets – a process called *reverse metabolomics* (**Fig. 1a**)^4,5^. Reverse metabolomics involves searching MS/MS spectra across large-scale LC-MS/MS data repositories to identify their occurrence in organisms, organs, health conditions, environments, or other metadata associations available with public data.

**Figure 1.**
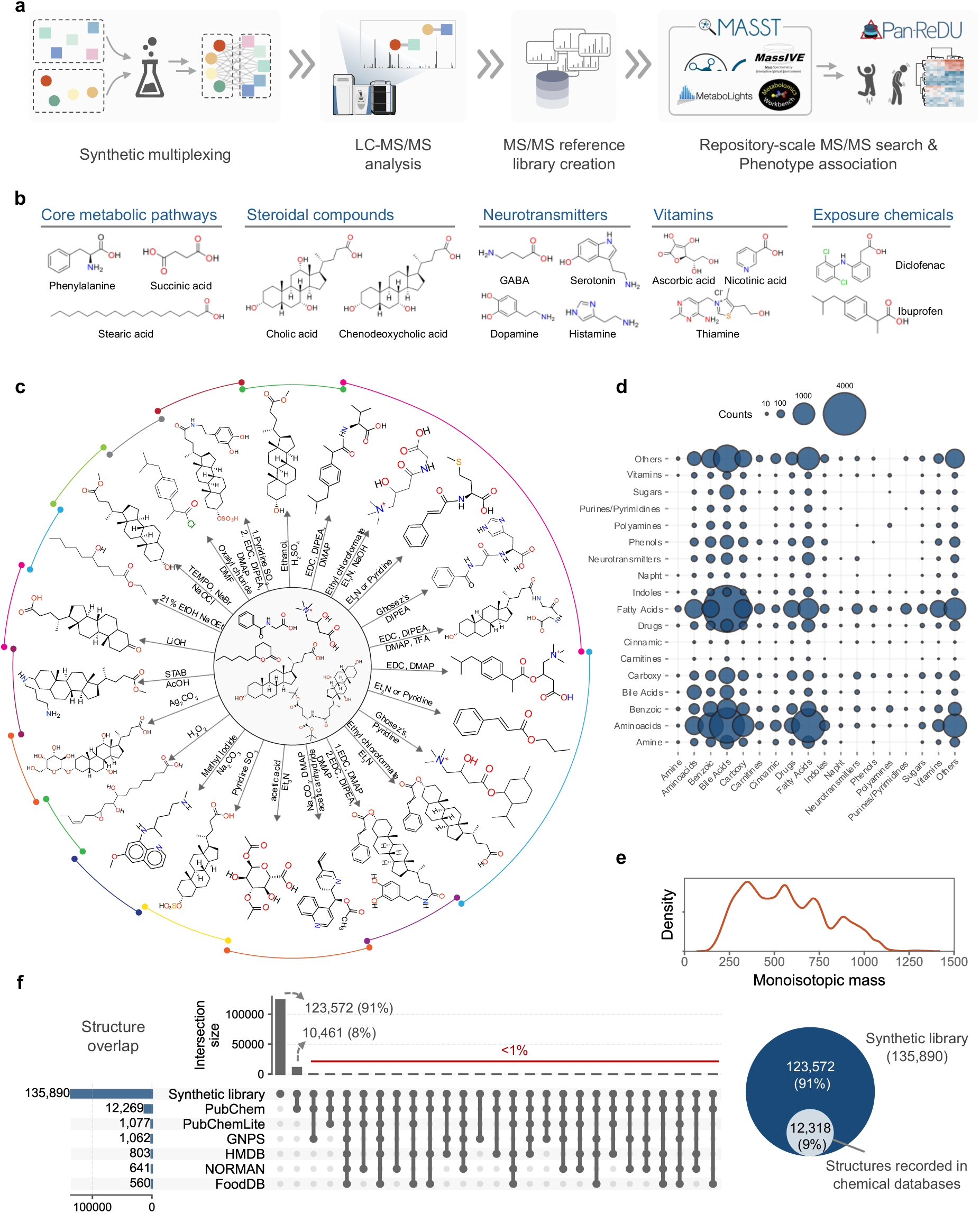
The creation of the multiplexed synthetic MS/MS library. **a)** Overview of the multiplexed synthesis-based reverse metabolomics^4,5^ performed in this work. Some of the products of reactions, such as acyl-chloride formation, result in intermediate reagents that subsequently undergo additional reactions. **b**) Representative molecules used as reagents in the multiplexed reactions. **c**) The types of reactions carried out in multiplexed reactions. **d**) Representation of unique structures present in the synthesis output (n=42,697). Chemical class categories include related molecules and derivatives. **e**) Mass distribution of the compounds that are part of the MS/MS library. **f**) Evaluation of the uniqueness of the MS/MS library compared to other structural databases.

This work provides both an annotation library to the community and demonstrates a technological advancement for searching across the ever-growing volume of untargeted metabolomics data. Our earlier work demonstrated the feasibility of reverse metabolomics^4^ by synthesizing approximately 2,400 compounds derived from 125 starting materials, including amino acids, bile acids, and lipids^5^. Searching the MS/MS spectra of these molecules across publicly available studies in the GNPS/MassIVE repository^6^ at the time resulted in new annotations of approximately 600 molecules. However, the scale of these searches at the time challenged the computational infrastructure, with some searches taking several days or weeks to complete. To enable searches at the scale of hundreds of thousands of MS/MS spectra – generated via multiplexed synthesis – we engineered a system capable of processing MS/MS search against >1.7 billion public MS/MS in milliseconds per query and thousands of queries per minute. This capability was enabled through a combination of hardware upgrades, algorithmic indexing strategies, and software engineering optimization.

The scale of MS/MS spectral comparisons required for this project required a dedicated expansion and engineering of the computational infrastructure capable of doing so. Reverse metabolomics analyses are now performed on a virtual machine equipped with two 64-core AMD processors and 2 TB of RAM, with public metabolomics data indices hosted on four SSDs to ensure rapid access. This setup supports high-speed spectral searches using indexed spectra – enabling the fast MASST (FASST) queries^7,8^. Due to the needs of this project, the second-generation GNPS platform^6^, the data and knowledge ecosystem that is being searched, has expanded to operate across five interconnected virtual machine servers: two equipped with dual 64-core AMD processors with 2 TB of RAM each, and three with dual 16-core CPUs totaling 768 GB of RAM. Data storage is distributed across two high-performance arrays, comprising 424 TB of SSDs – all linked through a 10 Gbit network backbone.

Together, this infrastructure underpins the GNPS2/MASST ecosystem, enabling community-scale reverse metabolomics and repository-wide MS/MS spectral searches at the necessary speed and depth. To broaden the search space, we indexed all data in GNPS/MassIVE^6^ and integrated additional public repositories, including MetaboLights^9^, Metabolomics Workbench^10^ and, more recently, NORMAN, a more environmentally focused repository, via the Pan-ReDU framework^11^. Searches can be performed using fast MASST^7^, along with its domain-specific variants (e.g., for microbes^12^, food^13^, plants^14^, tissues^15^, and microbiome-relevant studies^16^). In parallel, we enhanced the underlying data science by continuing to harmonize metadata vocabularies across these repositories, enabling MASST searches to return MS/MS spectral matches and additional relevant and interpretable metadata about the matched samples^11^. All indexed LC-MS/MS files, features, spectra, and synthetic reference libraries were converted to use Universal Spectrum Identifiers (USIs)^17^, ensuring complete provenance and traceability to the original deposited raw data. As of late August/ early September 2025, the number of LC-MS/MS files that are indexed and have harmonized metadata in PanReDU had grown to 920,790 LC-MS/MS files. This indexed infrastructure supports searches across 4,990 datasets from the repositories previously mentioned, comprising a total of 1,752,167,824 MS/MS spectra. These advancements eliminate previous computational bottlenecks and enhance the biological and environmental interpretability of reverse metabolomics results at this scale.

Using this infrastructure, we expanded the chemical space in this study by incorporating a more structurally diverse set of compounds representing a range of biologically and exposure-relevant starting materials that we leverage in multiplexed synthetic organic chemistry reactions. These included 1,450 small molecules that possessed functionality that was plausibly available for biotransformation in biological systems. The precursor molecules span core metabolic pathways (e.g., central carbon and fatty acid metabolism), steroidal scaffolds (e.g., bile acids), neurotransmitters, vitamins, dietary components, as well as exposure-related compounds such as dietary components, plastic-associated chemicals, ingredients from personal care products, chemicals used in manufacturing, and currently approved drugs (**Fig. 1b, Supplementary Table 1**). We prioritized compounds containing amines, carboxylic acids, and hydroxyl groups because their chemical reactivity in biological systems is well-characterized. These functional groups form esters, amides, oxidations, glycosylations, ethers and other common products, which can be readily generated in standard flask-based reactions. To model biologically and environmentally relevant biochemical transformations, we applied both single– and multi-step reactions with a multiplexed synthetic strategy, where multiple products are generated with multiple starting materials and reagents in one reaction vessel (**Fig. 1a**; details for each reaction can be found in **Supplementary Table 2**). These flask-based reactions were designed to emulate transformations commonly occurring *in vivo* or in the environment, including sulfation, conjugation, methylation, oxidation, hydrolysis, and amide formation (**Fig. 1c, d**). As the objective was to generate detectable products for MS/MS acquisition rather than maximize chemical yield, limited reaction optimization was carried out. No purification steps were needed given that we are using chromatographic separation, and due to the sensitivity of mass spectrometry, it is possible to detect and obtain MS/MS data for products with only a small amount of conversion in the multiplexed reactions. The mixtures with starting materials, reagents, and expected products were analyzed by LC-MS/MS using data-dependent acquisition.

To systematically annotate possible products from the LC-MS/MS data obtained from the multiplexed reactions, we developed a web application for *in silico* generation of all plausible structures of the products from our multiplexed and combinatorial reactions under defined conditions, as existing tools could not sufficiently scale. We then linked the MS/MS to ionic forms of the structures (with H^+^, Na^+^, NH_4_^+^, K^+^ adducts) that could be present in each synthetic reaction, including starting materials. This resulted in the 492,376 MS/MS spectra with molecular structure annotations that were synthesized. The remaining MS/MS that are not linked to structures are redundant MS/MS spectra of the same ion forms of molecules, other ion forms that we did not look for (e.g., in-source fragments, different adducts, and multimers)^1,2^, chimeric spectra, impurities, or unanticipated reactions. The outcome is an open and freely accessible MS/MS reference library. Due to redundant spectra for the same compounds, as well as isomeric overlap (as further discussed in the limitations section), the complete MS/MS library generated using multiplexing contains 172,483 candidate compounds represented by 134,453 unique MS/MS spectra, each indexed with a USI^17^. This means that when one searches with a USI in MASST and obtains a match to that particular MS/MS, one has to consider all isomers. The fact that, although more explicit for this library, one has to consider isomers is, in practice, similar to other MS/MS reference libraries based annotations (**see limitations discussion of this paper**). The candidate molecules that make up the multiplexed library cover diverse chemical classes relevant to both biological and environmental systems (**Fig. 1d**), spanning a mass range from approximately 150 Da to 1,350 Da (**Fig. 1e**). Given the synthesis prioritization of biologically relevant precursor molecules, we anticipated that a significant portion of this library would represent previously unexplored chemical space in biology. Based on the planar structures, 91% of these compounds were unique to our library and not present in any major structure databases (**Fig. 1f**). The highest overlap was obtained with PubChem^18^ (8%), which contains over 110 million structures. All other databases, including HMDB^19^, GNPS^6^, PubChemLite^20^, NORMAN^21^, and FooDB^22^, shared less than 1% overlap with the structures from the multiplexed synthetic MS/MS library (**Fig. 1f, Supplementary Table 3**).

Using the indexed FASST implementation, we searched the newly created MS/MS reference library against the public datasets (**Fig. 2a**). FASST was performed using ≥0.7 cosine score and ≥4 matching ions, criteria that typically result in an FDR <1%^23^. This search yielded matches to 60,146,352 indexed MS/MS spectra in pan-repository data. When combined with existing GNPS reference MS/MS libraries, a total of 8.1% of all MS/MS spectra across the indexed data across the repositories now have a library match, corresponding to a total annotation growth of 17.4%. Both the multiplexed synthesis library and existing libraries provide an initial structural hypothesis when a match within user-defined scoring criteria is obtained. Across all datasets, 63,369 MS/MS spectra from the multiplexed synthesis library were matched, corresponding to 15,190 distinct candidate structures (**Fig. 2b**). UMAP-based visualization of presence/absence patterns of the MS/MS across each taxonomic levels revealed that the MS/MS of many molecules were broadly distributed across many orders and other taxonomic levels (e.g., plants, fungi, animals), suggesting core or possibly even part of yet-to-be documented central metabolism, while others appeared to be taxon-specific (**Fig. 2c-d, Supplementary Figures 1-6**). MicrobeMASST^12^, which enables MS/MS searches against ∼60,000 LC-MS/MS of taxonomically defined microbial monocultures, revealed that 24,997 MS/MS spectra of synthesized compounds matched. After removing any candidate compounds that also matched to cultured human cells, this represents 4,596 candidate structures, or some related structural isomer, of putative microbial origin. Based on taxonomic information, most of the MS/MS that matched to cultured data from the bacterial phyla belonging to Actinomycetota, Pseudomonodata, Bacteroidota, and to a lesser degree Bacillota (**Fig. 2e**). In addition, we see matches to different fungi such as the Ascomycota and Basidiomycota phyla (**Fig. 2e**). This highlights that there is a large number of microbial molecules that can be readily accessed through synthesis that await to be fully explored. It should however be noted that the prevalence of the frequency is biased by the number of samples and conditions of LC-MS/MS files for a given taxonomic assignment available in the public domain. Based on NPClassifier^24^ classifications, molecules derived from alkaloids, fatty acids and terpenoids had their largest share of matches.

**Figure 2.**
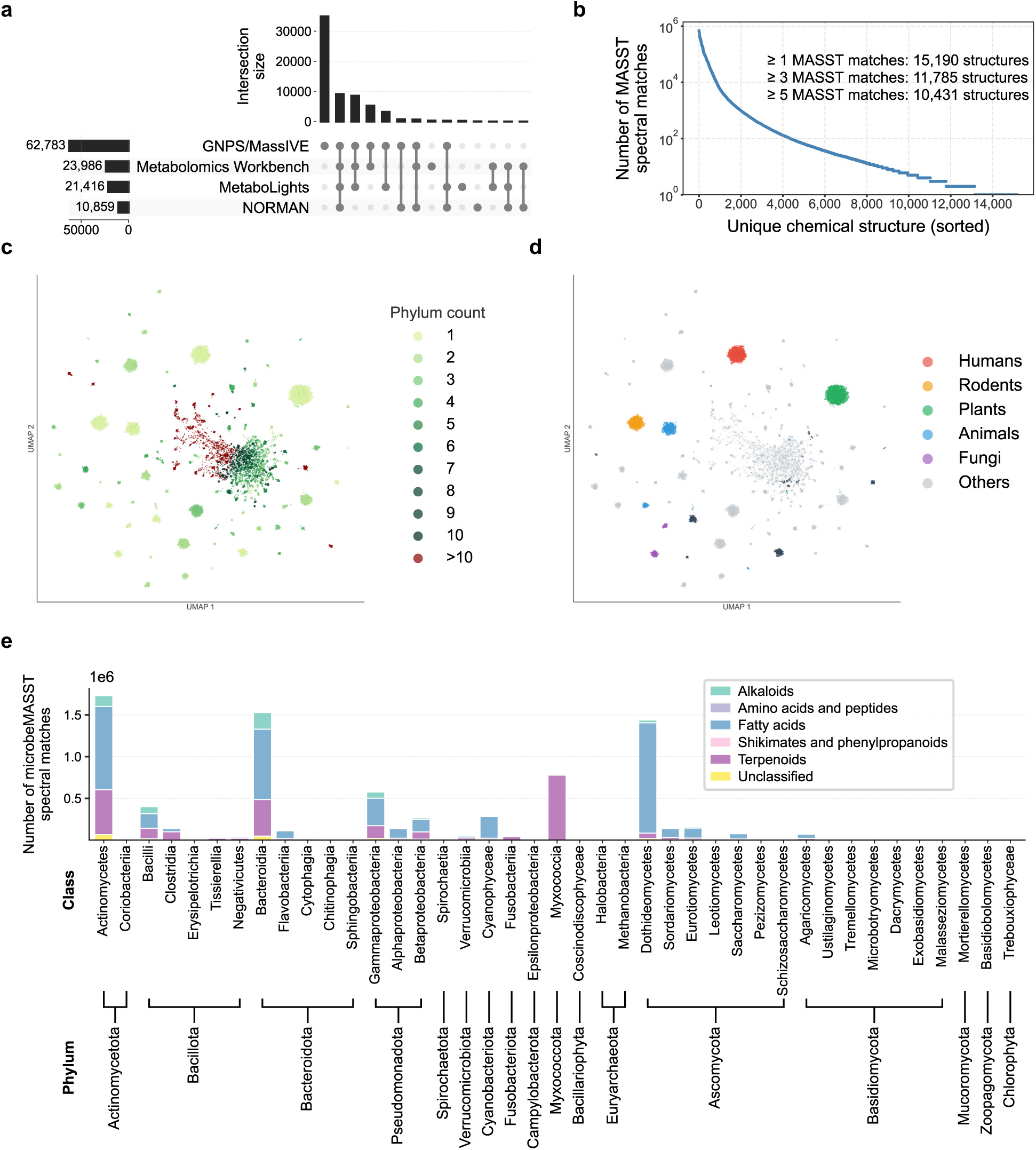
Large-scale reverse metabolomics across public datasets using the multiplexed synthetic MS/MS library. **a)** Upset plot showing the number of unique MS/MS spectra matched from the synthetic library to public datasets in GNPS/MassIVE, MetaboLights, Metabolomics Workbench, and NORMAN. **b)** Number of MASST spectral matches of the synthetic library for unique chemical structures. **c-d)** UMAP visualization of the taxonomic composition for a given MS/MS spectrum from the multiplexed library at the phylum level to which we had matched the MS/MS spectra across public datasets, highlighting both taxon-specific and widely shared metabolite signals. Each dot in the UMAP is an MS/MS spectrum from the multiplex library. UMAP of other taxonomic levels can be found in **Supplementary Figures 1-6. e)** Number of microbeMASST spectral matches of the synthetic library across microbial classes and phyla.

Of the 27,807 MS/MS spectra from the multiplexed synthetic library that matched *Homo sapiens* datasets, 2,679 were exclusive to human data (**Fig. 2d**), representing 1,404 candidate structures. Of these spectra, 6.0% (n=161) were derivatives of drug molecules. Examples include derivatives of ibuprofen, 5-aminosalicylic acid (5-ASA), atorvastatin, atenolol, primaquine, naproxen, and methocarbamol (**Supplementary Table 4**). Others include bile acids and their derivatives, fatty amides, peptides, carbohydrates, shikimates, phenylpropanoids, and alkaloid molecules. That we see matches to MS/MS generated from multiplexed reactions with drugs to human data only makes sense as, generally, other organisms (animals, including rodents, microbes, and plants) are generally not given these specific pharmaceutical compounds in the experiments that led to the generation of the untargeted metabolomics data available in the public domain. These spectra associated with humans were distributed across multiple body sites (**Fig. 3a**), with fecal samples showing the highest prevalence. Molecular networking of compounds detected exclusively in human samples revealed candidate drug-related metabolites, including MS/MS matches to 56 and 41 ibuprofen and of 5-ASA conjugates, respectively (**Fig. 3b**), the majority of which have not been previously reported. We obtained MS/MS matches corresponding to 29 5-ASA derivatives and 33 ibuprofen derivatives across 453,005 human LC-MS/MS datasets in Pan-ReDU (September 2025, **Fig. 3c-h**). MS/MS matches corresponding to 5-ASA derivatives were predominantly detected in human fecal datasets, whereas ibuprofen conjugate spectra were most frequently observed in human urine (**Fig. 3c,d**). Representative MS/MS matches are shown in **Fig 3e-h** and all others can be found as **Supplementary Figure 7**).

**Figure 3.**
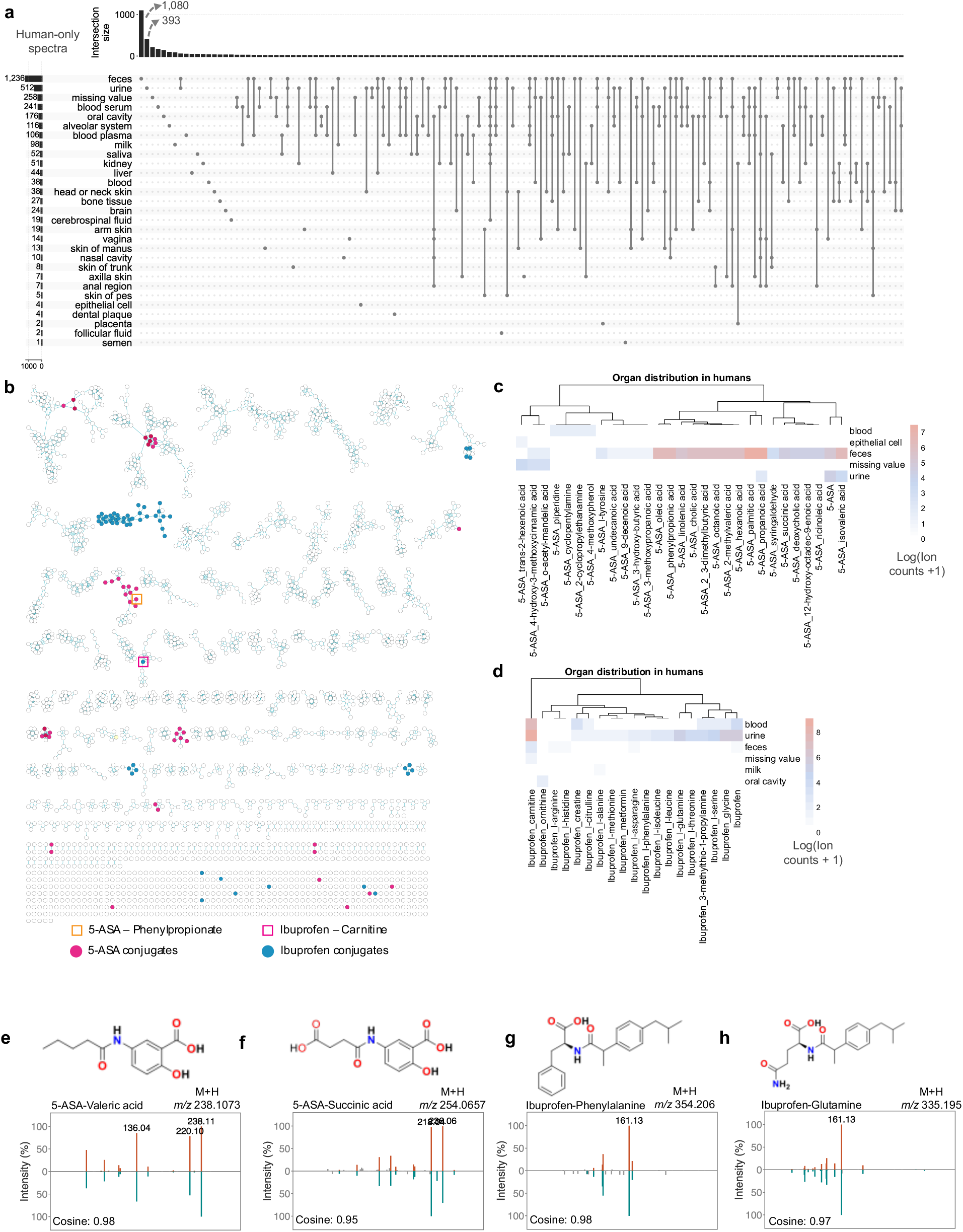
Molecules from the synthetic MS/MS library matched to human only data with MASST. **a)** UpSet plot showing the number of matched compounds associated with each parent drug and their overlaps across drugs. **b)** Molecular network of all matched compounds, with each parent drug and its associated matched derivatives colored distinctly. Two example clusters are highlighted: 5-aminosalicylic acid (5-ASA) and ibuprofen. **c–d)** Organ-level distribution of all matched derivatives for (**c**) 5-ASA and (**d**) ibuprofen across available human datasets, showing where these compounds were detected. **e–h)** Representative MS/MS mirror plots and chemical structures for selected matched analogs of 5-ASA and ibuprofen. Each plot displays the experimental MS/MS spectrum from the synthetic library (top) and the matched human spectrum (bottom), along with the corresponding compound structure. Full mirror plots and structures for all 5-ASA and ibuprofen derivatives are available in the Supplementary Information.

Although we used typical scoring conditions of cosine of 0.7 or higher that typically lead to less than 1% FDR for MS/MS spectral alignment^23^, MS/MS matches to reference libraries are always considered a structural hypothesis rather than a confirmed structural entity. We therefore set out to provide additional experimental validation of the existence of the 5-ASA and ibuprofen-derived annotations. In order to provide additional experimental support for the existence of these drug-derived metabolites, as they could also be derived from the post-chromatography electrospray droplet-based chemistry^25^, we would need to validate them with retention time and/or drift time in human samples against the synthesized standards. To accomplish this, given that our matches thus far were limited to public datasets, we used MASST to identify public-domain samples containing MS/MS spectra matching 5-ASA conjugates that we could match our standards against. Matches were found from inflammatory bowel disease (IBD) fecal datasets that we were able to get access to, enabling confirmation with MS/MS, retention time, and ion mobility using two different LC-MS/MS instruments (**Fig. 4a-c**). Across these fecal samples, we matched seven 5-ASA conjugates against synthetic standards, including four long-chain fatty acid conjugates, two short-chain fatty acid conjugates, and a phenylpropionate conjugate (**Fig. 4b**). Short-chain fatty acid and bile acid conjugates of 5-ASA have been previously reported as microbial metabolism products^26–28^, whereas the remaining conjugates have not been described before.

**Figure 4.**
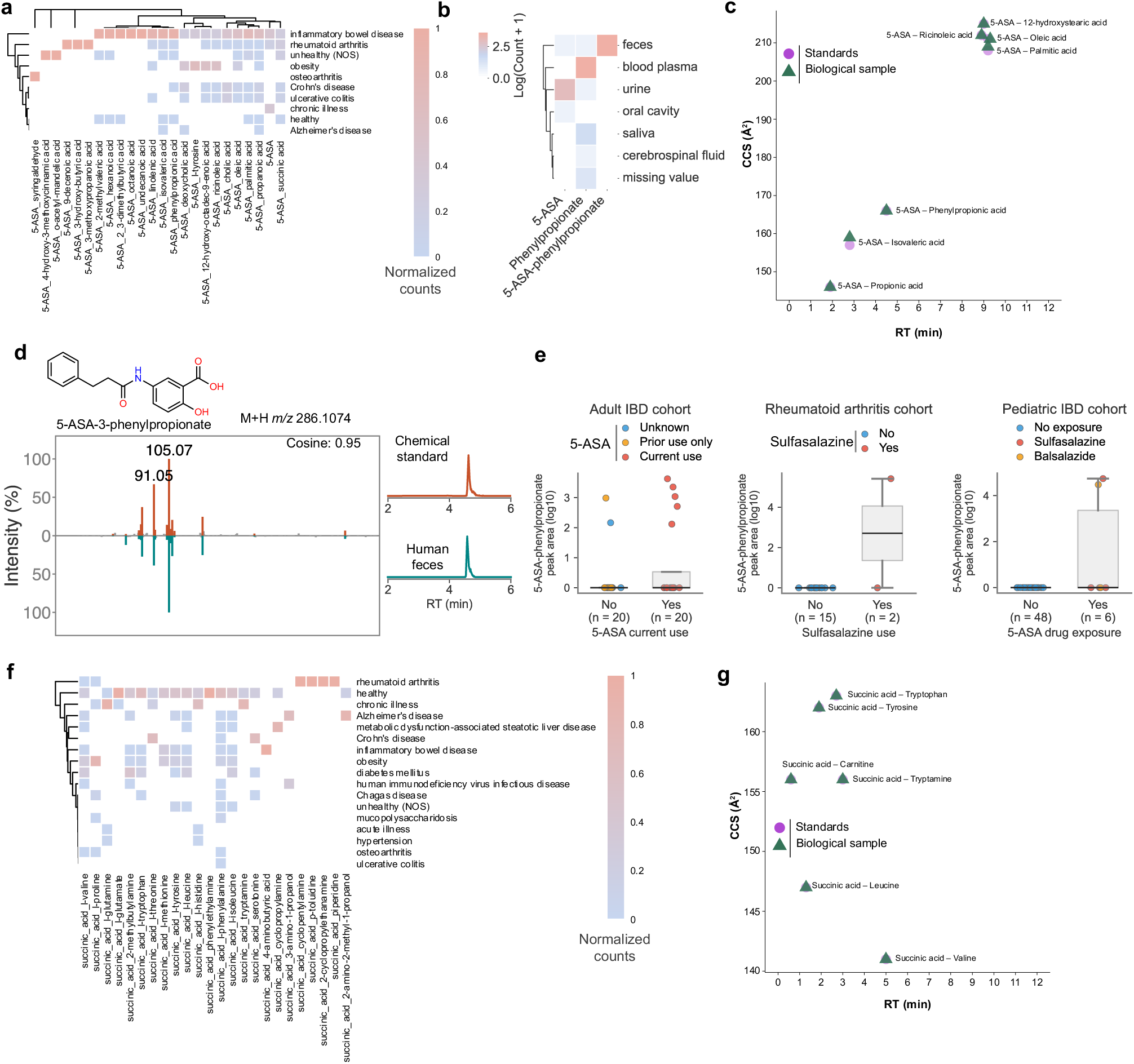
Characterization of 5-aminosalicylic acid (5-ASA) derivatives in human data. **a)** Association of 5-ASA and its derivatives with specific health conditions, highlighting links to IBD and RA. **b)** Comparative organ-level distribution of 5-ASA-phenylpropionate and its unconjugated counterpart phenylpropionate across human datasets. **c)** Experimental validation of 5-ASA derivatives using retention time (RT) and collision cross section (CCS) measurements, confirming MS/MS-based annotations (full MS/MS spectra in Supplementary Information). **d)** Representative MS/MS mirror plot and RT validation for 5-ASA-phenylpropionate, showing the synthetic spectrum (top) and matched human spectrum (bottom), alongside the compound structure. **e)** Log transformed peak areas of 5-ASA derivatives across two independent IBD and RA studies, illustrating medication based differences. **f)** Comparison of succinylated derivatives of 5-ASA across health conditions, demonstrating additional patterns of disease relevance. **g)** CCS and RT matches of succinic acid conjugates to chemical standards.

Focusing on the phenylpropionic acid, a known microbial metabolite^29^, we detected both 5-ASA and its phenylpropionic acid conjugate in feces (**Fig. 4c,d**). This conjugate was quantified in a fecal sample to be 20.8 µM (quantification details are available in the methods). Using MicrobiomeMASST^16^, a tool which links metabolites to microbiome-relevant information such as organs, age, interventions, and health conditions, 5-ASA-phenylpropionate was detected in 41/693 data files labeled as IBD, 6/333 data files labeled with rheumatoid arthritis (RA), and 1/14,567 healthy human samples^16^. Compared to data from healthy individuals samples, detection was more frequent in IBD (odds ratio [OR] = 916, 95% CI = 126–6669, Fisher’s exact p = 1.1 × 10⁻⁵⁴) and RA (OR = 267, 95% CI = 32–2226, p = 8.2 × 10⁻¹⁰). Direct comparison between IBD and RA revealed higher odds in IBD (OR = 3.43, 95% CI = 1.44–8.16, p = 0.0023), consistent with the more widespread clinical use of 5-ASA in IBD compared to RA. The single positive control labeled as healthy, we hypothesize was labeled incorrectly in the public domain data. It was part of a contrasting Western group in a microbiome study of remote villages^30^ and we hypothesized that this sample information to have been misassigned as clinical status, which was likely assumed to be healthy by the data depositor as it was a control group for a non-clinical study. Its inclusion therefore provides conservative effect size estimates, as exclusion would only further increase the odds ratios and strengthen the associations. Thus, while wide confidence intervals reflect uncertainty due to the rarity of the 5-ASA-phenylpropionate, enrichment in IBD and RA is robust, and the true effect is likely underestimated in our reported values.

As 5-ASA treatment is not universal among patients with IBD or RA, we next compared the observed metabolite detections to clinical metadata documenting 5-ASA or related prodrug administration, where available (**Supplementary Figure 8**). In three studies – two IBD cohorts and one RA cohort – with documented 5-ASA or prodrug usage (e.g., sulfasalazine, balsalazide), we compared 5-ASA-phenylpropionate matches to medication metadata. In a pediatric IBD cohort, 2 of 6 exposed study participants had matches, while 0 of 46 unexposed individuals did (**Fig 4e**). Combining all three datasets, structure matches were observed in 9/29 exposed, and 1/83 of non-exposed had detection. This supports that they were enriched in exposed individuals compared to unexposed controls (OR = 36.9, 95% CI 4.4 – 308.3, one-tailed Fisher’s exact test, *p* = 9 × 10⁻^5^), further supporting a drug origin for the phenylpropionate-linked 5-ASA compound consistent with known prescription patterns. In addition, it was observed in the monocultures of *Bacteroides caccae*, *Bacteroides vulgatus*, *Bacteroides luhongzhouii, Bacteroides thetaiotaomicron*, and in a 12-member Crohn’s disease synthetic community, to which phenylpropionic acid and 5-ASA were added to the growth culture. Thus confirming the microbial origin of the drug conjugate **(Supplementary Figure 8)**.

To confirm that 5-ASA conjugates reflect drug-specific exposure rather than baseline metabolic processes, we compared them to succinylated amines, metabolites expected to occur broadly. Succinic acid, a core intermediate of primary metabolism, reacts with diverse amines and is widely distributed across organisms and health states^31–33^. Indeed, Pan-ReDU yielded MS/MS matches to 26 succinylated amines (**Fig. 4f,g**), spanning humans, microbes, and other organisms (**Supplementary Figure 9**), with a large portion of the human matches (48.5%, n=928/1913) observed in healthy-labeled datasets. In contrast to the disease– and medication-restricted distribution of 5-ASA conjugates, succinylated amines were broadly detected, including in non-human data, highlighting that 5-ASA derivatives serve as specific markers of drug exposure rather than baseline metabolic products.

In contrast to succinylated amines and more similar to 5-ASA derivatives, the MS/MS matches to the 33 candidate ibuprofen-derived conjugates were found exclusively in human datasets, including data from healthy individuals (**Fig. 5a**). The many diabetes matches could be primarily driven by the preponderance of urine data for this health condition that are not as prevalent in other health conditions in the public domain data. Thus, although the biology may warrant further exploration, this observation could also reflect the database composition. Seven of these metabolites were verified by MS/MS and retention time across two different instrument platforms, one of which also provided ion mobility data, from human urine (**Fig. 5b-d**). In contrast to reported phase I/II transformations such as hydroxylation, carboxylation, glucuronidation, and taurine addition, we found no reported evidence for these conjugates in rodents or humans that were exposed to ibuprofen^34–37^. Instead, the ibuprofen-carnitine conjugate was widely observed across human datasets and across health categories **(Supplementary Figure 8)**, including healthy individuals – consistent with common over the counter use for muscle aches and headaches of individuals who would generally consider themselves healthy. It was detected in 105 human datasets. 61 files of which had available harmonized sample information, showing it spans feces, urine and serum/plasma, with urine being the most common matrix (**Fig. 5c**). Quantification in a urine sample gave a concentration of 1.4 µM. The presence of the conjugate in urine prompted us to test its consistency across samples with likely ibuprofen exposure. Analysis of fresh urine from an ongoing urogenital microbiome and metabolomic profiling study was performed to assess if the carnitine metabolite (and other ibuprofen-derived conjugates) were present. Urine samples were obtained from hospitalized patients likely to be receiving ibuprofen as part of their clinical care. The ibuprofen–carnitine conjugate was detected in all nine samples that also contained ibuprofen (**Fig. 5e**), and it was the most abundant ibuprofen-derived metabolite detected in all but one case, where the carboxy-ibuprofen signal dominated. Importantly, the discovery of these NSAID–carnitine conjugates was enabled by the multiplexed MS/MS library, which allowed structural annotation of metabolites that have eluded conventional metabolomics reference libraries. These libraries are largely restricted to commercially available standards and therefore rarely capture previously undescribed metabolites – despite ibuprofen having been approved by the FDA in 1974 and available over the counter since 1984. Because this is the first report of such carnitine conjugates, their potential biological consequences remain unknown. We therefore used this observation as a proof-of-principle that multiplexed synthesis can uncover biologically relevant molecules and set out to investigate the potential physiological role of ibuprofen–carnitine.

**Figure 5.**
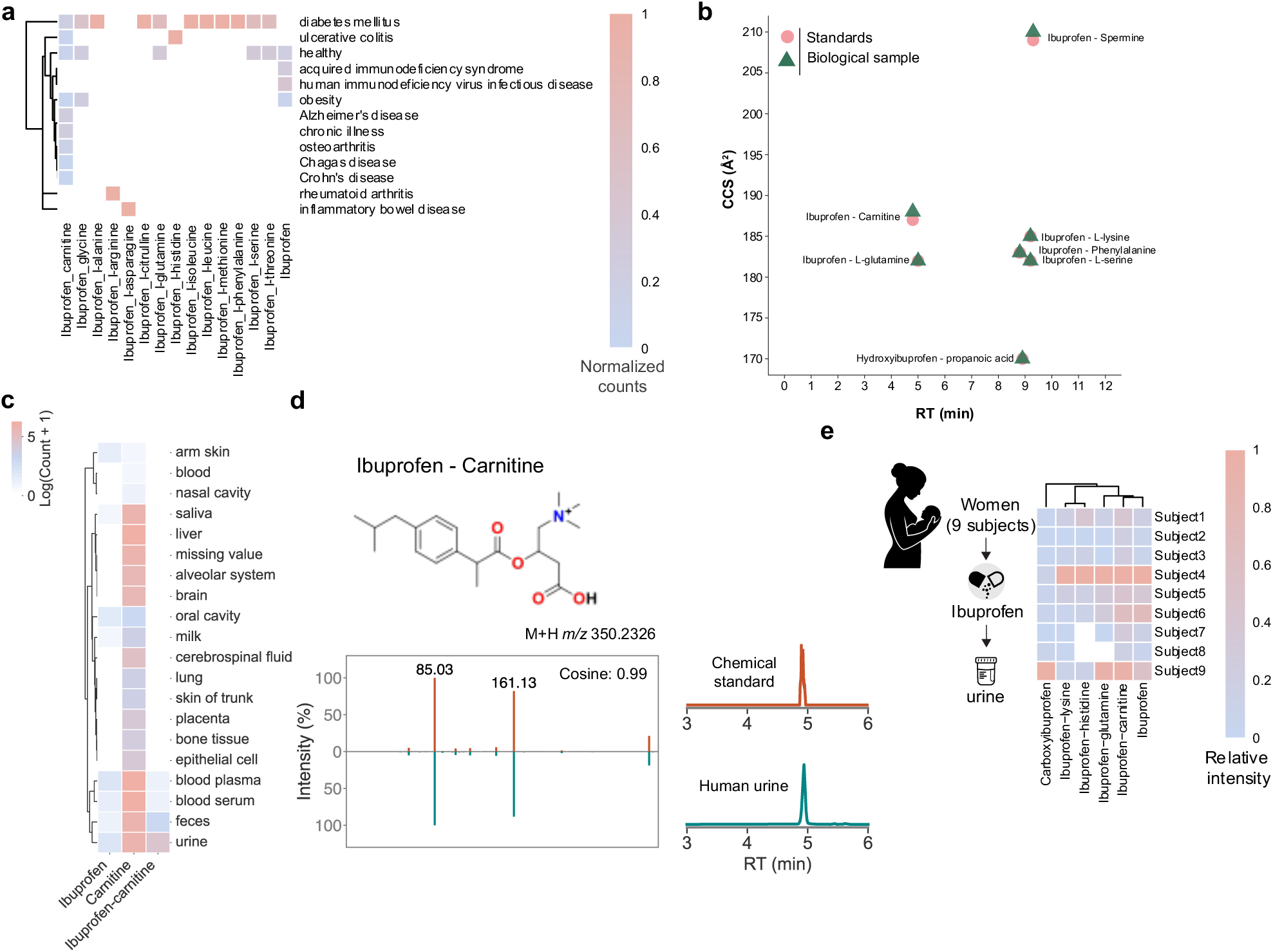
Characterization of ibuprofen-derived metabolites in human datasets. **a)** Distribution of the MS/MS of the ibuprofen parent compound and its conjugated derivatives across health conditions, highlighting associations with medication exposure and relevant disease groups. Each derivative is labeled by chemical class (e.g., carnitine, glucuronide, sulfate, acyl-conjugates). **b)** Experimental validation of ibuprofen derivatives using retention time (RT) and collision cross section (CCS) measurements; MS/MS spectra supporting annotations are provided in the Supplementary Information (or in the main figure, if included). Reported RT and CCS values are from authentic synthetic standards. **c)** Organ-level distribution map for IB-carnitine (representative ibuprofen-carnitine conjugate) showing repository-wide detection across tissues and biofluids; heatmap/points indicate the presence and relative frequency of matches in each organ dataset. **d)** Representative MS/MS mirror plot and RT validation for Ibuprofen-carnitine: synthetic library spectrum (top) versus matched human spectrum (bottom), with the annotated chemical structure and reported RT/CCS concordance. **e)** Distribution of all detected ibuprofen analogs in a hospitalized cohort demonstrating observed analog classes in urine. Heatmap colors represent the calculated peak area per detected molecule per subject divided by the maximum peak area of detected molecule across all subjects.

Ibuprofen, a drug commonly used to treat pain, inflammation and fever, has been described to decrease fatty acid β-oxidation by mitochondria^38,39^. β-oxidation is a major ATP-generating pathway and is critical for muscle function and regeneration after injury, and it requires carnitine for mitochondrial fatty acid transport. Case reports and clinical studies indicate that under conditions of high carnitine demand, such as muscle repair following endurance exercise or tissue remodeling in the postpartum period, impaired fatty acid oxidation can lead to muscle toxicity^40–47^. Endurance athletes have elevated ATP requirements and therefore increased reliance on carnitine-dependent lipid oxidation during recovery^43–46^, whereas pregnancy and postpartum states are associated with reduced systemic carnitine levels^47^. Moreover, individuals with genetic defects in carnitine transport may be vulnerable, consistent with clinical reports linking ibuprofen exposure to rhabdomyolysis in this population^40–42^. Prior to our discovery of ibuprofen-carnitine in humans, some NSAID-carnitine conjugates had been explored synthetically (e.g., naproxen– and ketoprofen-carnitine derivatives but not ibuprofen), motivated by the potential to enhance cellular uptake of the drug, via conjugation to carnitine, by the OCTN2 carnitine transporter and improve drug delivery to renal tissues while reducing systemic toxicity^47^. However, the occurrence of NSAIDs-carnitines *in vivo* in humans (or any other organism) had not been demonstrated. Interestingly, OCTN2 recognizes naproxen– and ketoprofen-carnitine derivatives as both substrates and inhibitors^48^. Thus, detection of ibuprofen–carnitine in human samples raises the possibility that such metabolites contribute to observed NSAID-related adverse effects, including mitochondrial or muscle toxicity, potentially through two non-mutually exclusive mechanisms: depletion of free carnitine and/or competition for carnitine transport into mitochondria.

We therefore asked whether ibuprofen–carnitine might interact with OCTN2, a key transporter responsible for carnitine uptake. Using Boltz-2 ensemble structural predictions^49^, co-folding models showed that ibuprofen–carnitine adopts a binding configuration similar to L-carnitine, forming three conserved hydrogen bonds, two with Arg471 and one with Tyr239, and additional π–cation interactions with Tyr358, Tyr239, and Phe443, consistent with the binding mode reported for other SLC22 family substrates^50,51^ (**Fig. 6a**). The model predicted for ibuprofen–carnitine that the ibuprofen moiety occupies an extended hydrophobic subpocket. Boltz-2 predicts that the affinity for ibuprofen-carnitine is nearly 50-fold better than carnitine. We tested if Ibuprofen-carnitine would interact with OCTN2 experimentally using a cell-based ¹⁴C-carnitine uptake assay in HEK293T cells stably expressing OCTN2-CeGFP and found that ibuprofen–carnitine reduced OCTN2-mediated carnitine transport (**Fig. 6b**)^52^. Uptake is expressed as a percentage of vehicle control after subtraction of background uptake in empty vector-transfected cells. Ibuprofen alone did not reduce OCTN2-mediated carnitine transport relative to vehicle (101.9 ± 1.2%), whereas ibuprofen–carnitine reduced carnitine uptake to 37.6 ± 2.3% of vehicle. Unlabelled carnitine served as a positive competitor control (19.3 ± 0.6%). To determine whether this reflects inhibition or reduction due to active but competitive transport, we quantified cellular carnitine and ibuprofen-carnitine after addition of carnitine or ibuprofen-carnitine to HEK293T cells containing a vector expressing OCTN2 where cells with an empty vector served as background. Supplementation with 5 µM and 50 µM carnitine produced a 2.2-fold and 5.9-fold increase in intracellular carnitine in OCTN2-expressing cells relative to empty vector controls, respectively, confirming active transporter-mediated uptake (**Fig. 6c**). When ibuprofen–carnitine was supplied as the substrate, OCTN2-expressing cells accumulated 64-fold and 47-fold more ibuprofen–carnitine than empty vector controls at 5 µM and 50 µM, respectively (**Fig. 6d**).

**Figure 6.**
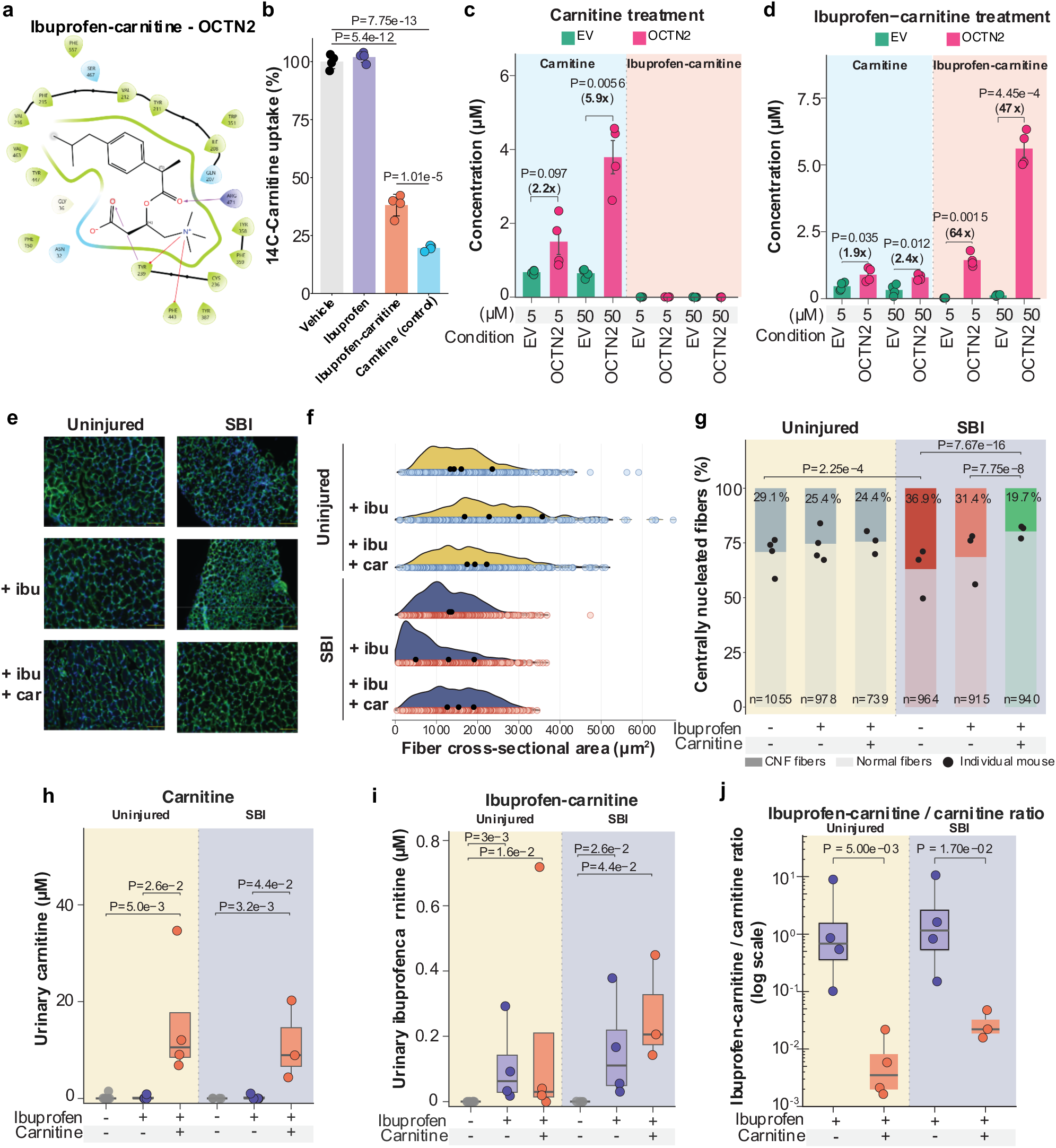
Biological and biochemical consequences of ibuprofen, carnitine, and ibuprofen-carnitine conjugate. **a)** Structural model of ibuprofen–carnitine bound to the carnitine transporter OCTN2, generated using Boltz-2 ensemble co-folding predictions, showing a binding configuration similar to L-carnitine with three conserved hydrogen bonds (two with Arg471, one with Tyr239) and π–cation interactions with Tyr358, Tyr239, and Phe443. **b)** Cell-based ¹⁴C-carnitine uptake in HEK293T cells stably expressing OCTN2-CeGFP, treated with ibuprofen, ibuprofen-carnitine, or unlabelled carnitine (positive competitor control). Uptake is expressed as a percentage of vehicle control mean (OCTN2 cells, no inhibitor). Data are shown as mean ± SD of individual replicates. Normality was assessed with the Shapiro-Wilk test and homogeneity of variance with Levene’s test: both assumptions were met. Group means were compared by one-way ANOVA followed by Tukey’s post-hoc test. **c–d)** Intracellular accumulation of **(c)** carnitine and **(d)** ibuprofen–carnitine in OCTN2-expressing or empty vector (EV) cells following incubation with 5 µM or 50 µM substrate. Bars show mean ± SEM with individual replicates. Pairwise comparisons between empty vector (EV) and OCTN2 within each substrate concentration were performed by Welch’s t-test; exact P values are shown. **e)** Representative fluorescence images of pubocaudalis muscle cross-sections immunostained for laminin (green) and DAPI (blue) from uninjured and simulated birth injury (SBI) mice treated with ibuprofen (+ibu) or ibuprofen plus carnitine (+ibu +car), acquired 14 days post-injury. All images are shown at the same scale. Scale bars are 100 µm in length. **f)** Density plot of the distribution of all measured muscle fiber cross-sectional area (µm²) across the treatment groups quantified from laminin-stained cross-sections. Individual data points indicate per-animal mean values. **g)** Percentage of centrally nucleated fibers (CNF) per animal across treatment groups in uninjured and SBI conditions. CNF remained elevated in ibuprofen-only treated injured mice, while co-treatment with ibuprofen and carnitine reduced CNF by ∼47% relative to the injured-only group and ∼37% relative to ibuprofen alone, indicating that carnitine promoted resolution rather than merely initiation of regeneration. No significant differences were observed among uninjured groups. Numbers within bars indicate group mean % CNF; numbers below bars indicate total fibers analyzed. Statistical comparisons were performed at the fiber level using Krustal-Wallis test followed by Dunn’s post-hoc test with Benjamini-Hochberg correction (BH); P values are indicated above the bracket. **h)** Absolute urinary carnitine concentrations in ibuprofen-exposed uninjured and SBI mice with and without carnitine supplementation. **i)** Absolute urinary ibuprofen–carnitine concentrations in the same animals as **(h)**. For **(h–i)**, each data point represents one biological replicate; boxes indicate median and interquartile range (IQR); whiskers extend to 1.5xIQR. Statistical comparisons were performed separately within uninjured and SBI groups using Wilcoxon test; raw P values are shown for significant comparisons. **j)** Urinary ibuprofen–carnitine:carnitine ratio across treatment conditions plotted on a log₁₀ scale, showing that carnitine supplementation markedly decreased this ratio in both uninjured and SBI animals. Groups were compared using Welch’s t-test on log-transformed ratios. Raw P values are shown. EV, empty vector; OCTN2, organic cation/carnitine transporter 2; SBI, simulated birth injury; CNF, centrally nucleated fibers; ibu, ibuprofen; car, carnitine.

To determine whether ibuprofen impairs muscle repair after injury as previously reported^47^, and to assess whether carnitine supplementation could rescue this phenotype, we used a balloon-induced birth injury mouse model. Ibuprofen-treated mice exhibited impaired muscle repair which was rescued by carnitine supplementation, directly linking carnitine availability to ibuprofen-associated defects in tissue regeneration. Histologic analysis of the pelvic floor muscle, pubocaudalis, 14 days after simulated birth injury confirmed that ibuprofen delayed muscle recovery, as reflected by reduced muscle fiber cross-sectional area (**Fig. 6e,f**), consistent with prior reports in rats^47^. We further show that carnitine supplementation restored tissue to levels comparable to the uninjured control, as evidenced by the increase in muscle fiber cross-sectional area, consistent with a role for carnitine in mitigating ibuprofen-impaired recovery (**Fig. 6f**). Centrally nucleated fibers, a marker of active muscle regeneration, were elevated following injury, as expected, and remained similarly elevated in injured mice treated with ibuprofen alone, indicating that regeneration was ongoing but incomplete. Notably, the ibuprofen and carnitine co-treated group showed a reduction in centrally nucleated fibers of nearly half (47%) compared to the injured only group, and 37% reduction to ibuprofen group only, suggesting that carnitine promoted the resolution of regeneration rather than merely its initiation, consistent with restored muscle repair (**Fig. 6f,g**). No significant differences were observed in centrally nucleated fibers among uninjured groups treated with ibuprofen or ibuprofen plus carnitine, despite a modest (16%) reduction trend, indicating that these effects are specific to the injured context.

Given that ibuprofen–carnitine conjugates can impair carnitine transport by OCTN2, and that ibuprofen impairs muscle repair in a manner that can be rescued by carnitine supplementation, we next asked whether impaired repair reflects depletion of free carnitine or altered ratios of conjugated to free carnitine. To illustrate why this ratio may be biologically relevant, consider an analogy: imagine a five lane highway leading into a two lane tunnel entrance shared by cars and buses, where cars represent carnitine and buses represent ibuprofen–carnitine. If the proportion of buses increases, fewer cars make it through the tunnel, even if the total number of cars outside the tunnel remains unchanged thereby affecting biology. Similarly, an increase in ibuprofen–carnitine could reduce the effective delivery of carnitine into cells by competing for OCTN2 transport capacity.

To examine both hypotheses, we performed quantification of carnitine and ibuprofen–carnitine in urine from ibuprofen-exposed mice, both injured and uninjured, with and without carnitine supplementation. Because carnitine is freely filtered by the kidney but efficiently reabsorbed in the proximal tubule through the high-affinity carnitine transporter OCTN2, urinary levels provide a systems-level readout of the balance between free and conjugated carnitine species interacting with this transporter. Carnitine levels were elevated in carnitine supplemented animals relative to unsupplemented controls in both non injured and injured conditions, while ibuprofen–carnitine was detectable in ibuprofen-treated animals but did not appear to increase in animals receiving carnitine supplementation (**Fig. 6h,i**). As the relative abundance of ibuprofen–carnitine remained relatively similar across conditions, and the amount of detectable carnitine increased by at least 64-fold in uninjured mice and 68-fold in injured mice upon carnitine supplementation, the ratio of ibuprofen–carnitine to carnitine decreased in animals receiving carnitine supplementation (**Fig. 6j**).

These observations support a model in which ibuprofen–carnitine competes with carnitine for OCTN2-mediated transport, reducing the effective intracellular availability of carnitine rather than depleting the systemic free pool. In this framework, which would require further detailed mechanistic studies, we hypothesize that increasing the proportion of conjugated species limits the amount of carnitine that can enter cells and become available to mitochondria for ATP production and muscle repair. We further hypothesize that carnitine supplementation mitigates this effect by increasing the free carnitine pool and diluting the conjugated fraction before carnitine and its conjugate are transported by OCTN2. Although demonstrated here in mice, future studies will be required to determine whether this mechanism operates in susceptible human populations and whether carnitine supplementation could prevent or ameliorate ibuprofen-associated muscle toxicity in humans.

Combined, these results demonstrate that reanalysis of public metabolomics data with a biochemically inspired multiplexed MS/MS library can uncover previously unrecognized metabolites that have biological consequences. In total, we detected putative MS/MS matches to 15,190 molecules in the public domain, of which 20 were elevated to level 1 identification according to the Metabolomics Standards Initiative through confirmation with MS/MS, retention time, and with ion mobility matching^10^. These annotations expand the known metabolic map, encompassing products of primary metabolism, host–microbe co-metabolism, and drug biotransformations. Importantly, the contrasting repository-scale distributions of 5-ASA conjugates, ibuprofen conjugates, and succinylated amines highlight how drug-derived metabolites – restricted to human data and reflective of medication use – can be distinguished from more broadly distributed primary metabolic conjugates. As it is not a traditional MS/MS library of compounds that can be purchased, we provide suggestions for its proper use as below.

### It is important to properly use and interpret the library and understand its limitations

Matches to the multiplexed MS/MS library, like any spectral reference in untargeted metabolomics, should be interpreted as plausible structural hypotheses rather than definitive identifications. Even high-scoring MS/MS library matches can be confounded by stereochemistry, positional or geometric isomerism, adduct variation, and fragmentation ambiguity. Tandem MS alone cannot generally unambiguously resolve single structure, so annotations must be evaluated in the context of biosynthetic logic, sample metadata, and orthogonal validation such as retention time, ion mobility, or isolation and NMR or other additional structural analysis. Assessing biological plausibility provides additional confidence. Harmonized metadata from public repositories enables evaluation across thousands of studies. For example, bile acids are animal-specific and their detection in plant datasets should be treated with caution, whereas very long-chain N-acyl lipids (>C24) are common in plants but rare in animals. Drug conjugates such as ibuprofen-carnitine or 5-ASA-phenylpropionate are observed only in human datasets where exposure is expected. Their absence in unrelated contexts further corroborates the annotation, while unexpected detections warrant deeper scrutiny. Similarly, co-occurrence of related metabolites provides additional evidence. Ibuprofen–carnitine often appears alongside other ibuprofen conjugates, and 5-ASA-phenylpropionate co-occurs with acetate, butyrate, and longer-chain fatty acid or amino acid 5-ASA conjugates, consistent with microbiome-mediated or host co-metabolism. In contrast, isolated detections may indicate rare transformations, false matches, or incomplete sampling. Integrating spectral evidence with co-occurrence and biochemical context helps translate MS/MS similarity into biologically meaningful hypotheses.

Tandem MS cannot reliably distinguish structural isomers based solely on fragmentation. Molecules with identical elemental composition, including those formed by acylation, amidation, or esterification, can yield similar spectra. For example, monoacetylation of OH’s of cholic acid produces three positional isomers whose MS/MS spectra are nearly indistinguishable under standard collision-induced dissociation. The multiplexed library prioritizes chemical diversity over site specificity, so many entries could represent mixtures or multiple isomers. Unambiguous structural assignment requires orthogonal validation, such as using retention time and drift time, as demonstrated for ibuprofen-, 5-ASA-, and carnitine-derived conjugates. Computational tools such as ICEBERG^53^ and Modifinder^54^ are expected to enhance isomer discrimination and annotation coverage in the future.

All spectra were acquired on a single instrument platform under defined collision-induced dissociation conditions, detecting primarily [M+H]^+^, [M+Na]^+^, and [M+NH_4_]^+^ adducts. Consequently, less common ion forms, multimers, or side products are underrepresented. The library spans over 4,000 reactions collected in positive ion mode, reflecting the majority of public LC-MS/MS data, though negative mode and additional instrument platforms would expand coverage. From ∼10 million spectra generated, ∼0.5 million were curated into the final library. The remaining spectra likely include uncharacterized analogs, unanticipated reactions, different ion forms, or in-source fragments. These data are publicly accessible for future reanalysis, reaction discovery, and iterative library expansion through molecular networking and annotation propagation.

Naming the compounds is a major challenge and we welcome anyone reading this to reach out for practical solutions. Many synthesized compounds are absent from structural databases, so conventional names are often impractical. IUPAC chemical names generated from SMILES are too complex for routine use. To improve clarity and computational accessibility, a reagent-based naming convention was adopted. For instance, we use the name “Erythro-aleuritic acid_glycine (known isomers: 0; isobaric peaks: 2)” This denotes a reaction between Erythro-aleuritic acid and glycine, with the underscore separating reagents and parentheses indicating predicted isomers and observed peaks. Each entry links to its raw file and one of the possible SMILES representations. While this system enhances traceability, it does not resolve inherent structural ambiguity. Matches should therefore be treated as biologically plausible leads, guiding hypothesis-driven synthesis and annotation refinement through orthogonal validation as done with 5-ASA and ibuprofen conjugates.

## Conclusion

This work establishes a scalable, hypothesis-driven framework for reverse metabolomics by combining biologically inspired multiplexed synthesis, high-resolution MS/MS, and systematic mining of public datasets. The resulting synthetic MS/MS reference library – comprising nearly half a million curated spectra across structurally diverse small molecules – enables broad exploration of previously unannotated chemical space. Through iterative workflows of match → hypothesis → synthesis → reanalysis, researchers can uncover unexpected biochemical transformations, such as those demonstrated with bile acids, N-acyl lipids, carnitine, carbohydrates and clinically relevant drug conjugates with ibuprofen and 5-ASA.

This approach is not static: it will evolve. A key long-term goal is to curate every MS/MS spectrum – irrespective of ion form, adduct, or fragmentation condition – so that each signal in LC-MS/MS based metabolomics data can eventually be linked to an interpretable structural hypothesis. Even when multiple ion forms, in-source fragments or droplet based chemistry get annotated, their inclusion enables researchers to make informed decisions about how to process, quantify, or exclude those signals depending on their biological relevance and analytical context^1,25^. As new hypotheses arise and uncharacterized MS/MS features accumulate, the system can be expanded to include additional compound classes – ranging from dietary and environmental exposures to microbiome– and host-derived metabolites. Future efforts should also incorporate more complex, multi-step, or enzyme based synthetic transformations to further mimic biochemical metabolism and extend annotation capacity into deeper regions of chemical space that are not yet being explored.

Our detection of ibuprofen–carnitine, together with new data showing that the conjugate inhibits the carnitine transporter, and that carnitine supplementation rescues delayed muscle recovery post-pregnancy injury in mice, supports a mechanistically grounded hypothesis consistent with both our results and prior literature. Ibuprofen has been linked to rhabdomyolysis in individuals with rare carnitine transport mutations, in high-demand physiological states (e.g., endurance exercise, pregnancy), and in overdose contexts, suggesting that NSAIDs may act as mitochondrial stressors in genetically or physiologically susceptible individuals. Although carnitine conjugation is documented for other carboxylic acid drugs (e.g., valproate, pivoxil-containing prodrugs)^55^ via acyl-CoA intermediates, often leading to carnitine depletion or toxicity, this pathway has not previously been described for ibuprofen despite its widespread use. Given that several NSAIDs form acyl-CoA thioesters, it is plausible that carnitine conjugation to NSAIDs is more common than recognized. Such conjugates could compete with endogenous fatty acylcarnitines for mitochondrial import, reduce free carnitine through urinary loss, and disrupt energy metabolism in high-demand tissues such as skeletal muscle. These findings support a hypothesis where a subset of the population may be at increased risk for NSAID-induced mitochondrial or muscle toxicity due to carnitine depletion or competition, particularly under chronic exposure to carboxylic acid–containing drugs.

Ibuprofen-carnitine would have evaded detection without the multiplexed synthesis MS/MS library and the ability to search against 100,000s of LC-MS/MS data files from human samples. While tandem MS has inherent limitations – including difficulty distinguishing structural isomers, variability in ion forms – these challenges are met with scalable data science strategies. Molecular networking and nearest-neighbor propagation allow for class-level annotations beyond exact matches. Mass difference analysis would be expected to link a significant portion of unmatched features to related molecules^56^ that did not yet make it in the 2025 multiplexed library. These would correspond to predictable modifications of curated structures^57^, emphasizing the opportunity to grow a future library.

Ultimately, this multiplexed synthesis strategy represents a unique route to illuminate the “dark matter” of the metabolome^2^. It facilitates data-driven structural hypothesis generation, structural anchoring of unknowns, and scalable annotation workflows that bridge synthetic chemistry, informatics, and biology. In doing so, it will also require shifts in the field from static pathway representations of hand curated metabolic pathway maps toward dynamic, interconnected computationally created metabolic networks that not only can handle annotation ambiguity but also reflect the true diversity and complexity of life’s chemistry – empowering future discoveries across metabolomics, exposomics, pharmacology, and systems biology.

## Methods

We have significantly expanded the reverse metabolomics approach, which associates MS/MS spectral profiles of the synthesized compounds with biological phenotypes through analysis of extensive public untargeted metabolomics repositories. We have extended its capacity in both computational capabilities and by creating a large reference spectral library to enable more comprehensive discovery of complex metabolites and its chemical–biological association. At its foundation, reverse metabolomics identifies the occurrence of any submitted MS/MS spectrum within public repositories and subsequently utilizes the associated metadata to correlate metabolites with a range of experimental variables, including disease states, taxonomic distribution, and sample types.

To validate the enhanced scalability of this methodology, we generated a library of structurally diverse candidate metabolites via multiplex synthesis and acquired corresponding LC-MS/MS data. Our multiplex synthesis encompassed five principal compound classes: amino acid conjugates (N-acyl amides, glutathione adducts, and peptide derivatives); microbial–host co-metabolites (bile acid amidates and bile acid esters), secondary metabolites (phenolic glycosides, alkaloids, and terpenoids); lipid classes and derivatives (fatty acid esters, glycerophospholipids, and sphingolipids); and xenobiotics (drug metabolites, environmental contaminants, and dietary compounds). The occurrence of the synthesized compounds in the public domain was obtained using Mass Spectrometry Search Tool (MASST)^7^, and the relevant metadata was analyzed and assessed using Reanalysis of Data User Interface (Pan-ReDU)^11^.

The multiplex synthetic MS/MS spectra exhibit a distribution of molecular ion forms, including 297,483 spectra (60.4%) corresponding to [M+H]⁺, 92,872 spectra (18.9%) to [M+NH_4_]⁺, and 53,269 spectra (10.8%) to dehydrated ions ([M+H–H_2_O]⁺). The remaining spectra represent less common adducts, such as [M+Na]⁺, [M+K]⁺, and doubly dehydrated ions ([M+H–2H_2_O]⁺).

## Chemical class prediction

The compound class information of newly synthesized chemicals were predicted using NPClassifier^24^, a deep neural network-based tool for structural classification. This was programmatically achieved via the GNPS2 API using the SMILES strings. For compounds which had more than one possible compound pathways provided by NPClassifier, the first pathway was reserved for downstream analysis and visualization.

## Sample collection and extraction

Urine samples were collected as part of an ongoing prospective cohort study for benchmarking storage and processing of the urogenital microbiome and metabolomic profiles (UC San Diego IRB#801735). Urine samples were obtained from hospitalized patients likely to be receiving ibuprofen as part of their clinical care. Voided urine was self-collected by participants and aliquoted and frozen at –80 until extraction. Urine samples were prepared as previously described^58^.

In brief, a 200 µL aliquot of urine was transferred into an empty sample tube. Then, 800 µL of 80% methanol was added, resulting in a final volume of 1 mL. Samples were vortexed for 5 s and then incubated at −20 °C for 20 min for protein precipitation. Following incubation, samples were centrifuged at 2000 rpm for 5 min at 4 °C to pellet the precipitated proteins. A volume of 800 µL of the resulting supernatant was transferred into the wells of a pre-labeled 96 deep-well plate. Samples were dried using a centrifugal vacuum concentrator (Centrivap). The dried residues were reconstituted in 250 µL of a 50% methanol-water solution containing 1 µM sulfadimethoxine as an internal standard. Fecal samples were obtained from a pilot study investigating the influence of diet on patients with rheumatoid arthritis (UC San Diego IRB#161474). Stool samples were weighted and extracted at a ratio of 50 mg of sample to 800 µL of 50% MeOH/H_2_O. A 5 mm stainless steel bead was added to the samples and homogenized using a Qiagen TissueLyser II for 5 min at 25 Hz, before being incubated overnight at 4 °C. Samples were centrifuged at 15,000 x g, 200 µL was transferred, and dried using a CentriVap. All samples were resuspended with 200 µL containing an internal standard and incubated at –20 °C overnight. All samples were centrifuged at 15,000 x g and 150 µL of supernatant was transferred into a glass vial for LC-MS/MS analysis.

## LC-MS/MS data collection

Biological samples and the synthetic standards were obtained for retention time and MS/MS spectral matching and were subjected to LC-MS/MS analyses. The LC-MS/MS analyses were carried out with a Vanquish UHPLC system coupled to a Q-Exactive Orbitrap mass spectrometer (Thermo Fisher Scientific, Bremen, Germany). The chromatographic separation was performed on a Polar C18 column (Kinetex C18, 100 x 2.1 mm, 2.6 μm particle size, 100A pore size – Phenomenex, Torrance, USA), and the mobile phase consisted of H_2_O (solvent A), and ACN (solvent B), both acidified with 0.1% formic acid. The following gradient was employed to evaluate retention time matching between synthetic standards and the compounds present in the samples: 0-0.5 min 5% B, 0.5-1.1 min 5-20% B, 1.1-5.0 min 20-40% B, 5.0-9.0 min 40-100% followed by a 1.5 min washout phase at 100% B, and a 1.5 min re-equilibration phase at 5% B. The flow rate was set at 0.5 mL/min, the injection volume was fixed at 3 μL, and the column temperature was set at 40 °C. Data-dependent acquisition (DDA) of MS/MS spectra was performed in the positive ionization mode. Electrospray ionization (ESI) parameters were set as: 52.5 AU sheath gas flow, 13.75 AU auxiliary gas flow, 2.7 AU spare gas flow, and 400 °C auxiliary gas temperature; the spray voltage was set to 3.5 kV and the inlet capillary to 320°C and 50 V S-lens level was applied. MS scan range was set to 300-800 *m/z* with a resolution of 35,000 with one micro-scan. The maximum ion injection time was set to 100 ms with an automated gain control (AGC) target of 1.0E6. Up to 5 MS/MS spectra per MS1 survey scan were recorded in DDA mode with a resolution of 17,500 with one micro-scan. The maximum ion injection time for MS/MS scans was set to 150 ms with an AGC target of 5E5 ions. The MS/MS precursor isolation window was set to 1 *m/z* with an offset of 0 *m/z*. The normalized collision energy was set to a stepwise increase from 25, 40, and 60 with *z* = 1 as the default charge state. MS/MS scans were triggered at the apex of chromatographic peaks within 2 to 5 s from their first occurrence. The quality and reproducibility of the analyses were evaluated considering the retention time and the *m/z* of a standard solution containing a mixture of six standards (amytriptiline, sulfamethizole, sulfamethoxine, sulfadimethoxine, coumarin 314, and sulfachlopyridazine) which was analyzed every five samples.

## MS/MS spectral library generation

### Structure generation of expected product molecules

To generate the structural information (SMILES strings) of the expected chemical products, we first create an input csv file containing compound names and SMILES strings of the reactants. This file is then uploaded to the AutoSMILES app. Next, we select which reactant to use as the sample ID, and specify the first and second reactant locations in the input csv file. Then, we enter the desired number of decimal places for mass precision. The type of reaction to perform is then selected – for example, amidation, esterification, hydroxylation, methylation, etc. Finally, the app generates all the product SMILES strings for the input reactants. The pipeline can be accessed via rxnSMILES · Streamlit.

## MS/MS spectra retrieval from raw LC-MS/MS data

Raw LC-MS/MS data collected for the multiplex synthesis library creation were first converted into an open format (mzML) using MSConvert. Then the mzML files and the csv files containing product SMILES strings were uploaded to the GNPS2 (GNPS2) file browser. The reverse_metabolomics_create_library_workflow was applied on the input mzML files and compound csv files, creating the MS/MS library in the format of mgf and tsv. These output mgf and tsv files were then uploaded to the GNPS library GNPS Library. The library generation workflow is available on GNPS2 through https://gnps2.org/workflowinput?workflowname=reverse_metabolomics_create_library_workflow.

## Repository-scale MASST searches

A minimum of 0.7 cosine score and 4 matched peaks was used to collect similar or identical MS/MS spectra for the four main metabolomics repositories (GNPS/MassIVE, Metabolomics Workbench, Metabolights, and NORMAN). Both MS1 and MS/MS mass tolerances were set to 0.05 Da. Any spectral match against the multiplex synthetic datasets deposited on GNPS/MassIVE were removed before analysis.

## Analysis of drug conjugates

Ibuprofen and 5-ASA conjugates were multiplex-synthesized and subjected to the reverse metabolomics workflow^4^. For the MASST analysis as shown in **Fig. 3d**, a minimum of 3 matched peaks was applied to account for small molecules such as 5-ASA and phenylpropionate which do not usually generate more than 3 peaks in their MS/MS spectra. As these drugs were only provided to humans, we removed any drug conjugates with MS/MS that matched to non-human samples, as shown in heatmaps in **Fig. 4b**. A total of 21 ibuprofen and 29 5-ASA conjugates were successfully identified. In addition to these, further matches were observed with other nonsteroidal anti-inflammatory drugs (NSAIDs), including aspirin, naproxen, and various NSAID metabolites. Notably, several matches were also detected with non-NSAID pharmaceuticals, such as atorvastatin, atenolol, metoclopramide, and primaquine, suggesting a broader interaction profile across multiple drug classes.

## Foundational model Boltz-2

The co-folded structures of L-carnitine, Ibuprofen–carnitine, and the organic cation/carnitine transporter 2 (OCTN2) protein were generated using the foundational model Boltz-2. All structure predictions were performed on an NVIDIA RTX 4090 GPU using the default Boltz-2 hyperparameters, specifically three recycling steps for prediction, 200 sampling steps during diffusion, and one diffusion sample per inference. Potentials-based terms were enabled to improve local geometries. For each inference, the SMILES representation of L-carnitine was obtained from PubChem (CID: 10917), and the SMILES representation of Ibuprofen–carnitine was produced by manually drawing the chemical structure in the PDB Chemical Sketch tool. The amino-acid sequence of OCTN2 was retrieved from UniProt (Entry Name: S22A5_HUMAN; Accession: O76082). The resulting co-folded structures were analyzed in ChimeraX (version 1.10rc202505280131)^59^ to identify ligand–protein hydrogen bonds using default geometric criteria, and these analyses were used to visualize the ligand-binding site. The structures were further examined in Maestro 14.6 (Schrödinger) to generate detailed images of ligand–protein interactions (cutoff of 4 Angstroms).

## Analysis of co-folding structures

Both co-folding structures—L-carnitine and ibuprofen-carnitine—exhibit three hydrogen bonds: two with Arg471 and one with Tyr239. Consistent with what has been reported for other SLC22 family members, the quaternary ammonium in both ligands is further stabilized through π–cation interactions with Tyr358, Tyr239, and Phe443.

However, the ibuprofen–carnitine conjugate engages the binding pocket more extensively. Its larger hydrophobic and aromatic moiety occupies a greater portion of the OCTN2 cavity, generating a broader hydrophobic enclosure involving multiple residues. This enhanced pocket filling allows the ligand to form additional stabilizing interactions, resulting in overall stronger binding to the transporter.

This trend is reflected in the Boltz-2 ensemble predicted IC₅₀ values of L-carnitine: 165.85 μM and Ibuprofen–carnitine: 3.4 μM. Although the predicted IC₅₀ for L-carnitine is much higher than the experimentally reported value of 4.3 μM^60^, the model still clearly captures the relative affinity shift, indicating that ibuprofen–carnitine is predicted to be a substantially stronger inhibitor of OCTN2.

## Uptake assay

The *in vitro* uptake assays were performed using methods previously described (PMID: 36343260). To determine the effect of ibuprofen-carnitine on OCTN2-mediated carnitine transport, HEK293T cells stably expressing OCTN2-CeGFP or empty vector were seeded at a density of 50,000 cells per well (100 µL) in poly-D-lysine-coated 96-well plates (PerkinElmer, Cat# 6005040) approximately 24 h prior to the uptake studies. The culture medium (DMEM supplemented with 10% FBS) was removed, and the cells were washed twice with Hank’s buffered salt solution (HBSS) pre-warmed to 37 °C. After the second wash, cells were incubated in HBSS for 10 minutes at 37 °C. Uptake was initiated by adding HBSS containing trace amounts of 14C-L-carnitine (Moravek Biochemicals, #MC1147) supplemented with either vehicle (0.1% DMSO), 100 µM unlabeled L-carnitine, 100 µM ibuprofen, or 100 µM ibuprofen–carnitine. After 15 minutes incubation at 37 °C, the uptake reactions were terminated by washing the cells two times with 200 µL ice-cold HBSS. Cells were then lysed by the addition of 100 µL MicroScint fluid (PerkinElmer) and incubated on a shaker for 1 h. Radioactivity was quantified using a liquid scintillation counter (MicroBeta2® microplate counter, PerkinElmer).

For ibuprofen-carnitine uptake, HEK293T cells stably expressing OCTN2-CeGFP or empty vector were seeded at a density of 200,000 cells per well (500 µL) in poly-D-lysine-coated 24 well plates and cultured for 24 h to reach 90-95% confluency. Prior to uptake studies, the culture medium was aspirated, and the cells were incubated in 0.5 mL of HBSS at 37 °C for 10-20 minutes. Uptake was initiated by adding ibuprofen-carnitine (5, 50 μM) or carnitine (5, 50 μM) in HBSS for 20 minutes at 37 °C. The uptake reactions were terminated by washing the cells twice with 0.8 mL of ice-cold HBSS buffer. Intracellular metabolites were extracted by incubating the cells in 400 μL of methanol for 30 min with shaking at room temperature. From each well, 300 µL of the methanol extract was transferred to a 1.5 mL microcentrifuge tube and stored at −80 °C until quantification via LC-MS/MS analysis.

## Carnitine and Ibuprofen-carnitine conjugate quantification in carnitine and ibuprofen exposed mice, injured and uninjured

A matrix-matched assay was developed for the absolute quantitation of carnitine and ibuprofen–carnitine in mouse urine samples and HEK293T cell extracts to evaluate uptake via the OCTN2 transporter. Carnitine and ibuprofen–carnitine standards were accurately weighed using an OHAUS™ 30100600/EMD microscale analytical balance (Fisher Scientific, USA) with a precision of 0.0001 g and dissolved in 1 mL methanol to prepare 1 mM stock solutions. Serial dilutions were performed to generate calibration standards over a concentration range of 0.001 nM to 10 µM. To achieve matrix-matched conditions that mimic biological urine, pooled urine was spiked into the calibration standards. A non-isotopic internal standard (sulfadimethoxine) was prepared in the same manner and added to all samples at a final concentration of 1 µM prior to LC–MS data acquisition, as described above. Calibration curves were constructed using peak area ratios derived from extracted ion chromatograms for carnitine and ibuprofen–carnitine (Skyline, version 21.2.0.425). Peak areas were background-corrected by subtracting signals from blank samples. A weighted second-degree polynomial regression model (inverse response weighting) was applied to generate calibration curves, and the resulting quadratic equations were used for quantification. The limits of detection (LOD) and quantification (LOQ) were calculated as LOD = 3.3 (σ/S) and LOQ = 10 (σ/S), where σ represents the standard deviation of the y-intercept and S represents the slope of the calibration curve. Precision was assessed using the coefficient of variation (CV), and accuracy was determined by comparing measured concentrations to theoretical values (n = 3 injections). All calculated concentrations were corrected for dilution factors to reflect physiological levels. Replicate injections (n ≥ 3 per sample) were used to determine CV values (**Supplementary Table 8**).

## Histological analysis of mice muscle tissue

Three-month-old C57/BL6 late-pregnant mice were randomly assigned to uninjured and birth injured groups at ∼embryonic day 17.5. Birth injured groups were subjected to vaginal balloon distension that replicates pelvic injury experienced during vaginal birth. Briefly, animals undergoing vaginal distension were anesthetized, a 6F transurethral catheter was inserted into the vagina and a balloon was inflated with 1 ml. A 13 g weight was attached to the end of the catheter for 1 hour to replicate the circumferential and downward strains associated with parturition. The day after birth injury, uninjured and birth injured groups were randomly assigned to no treatment, ibuprofen treatment, or ibuprofen+carnitine treatment, with group numbers as follows: Uninjured: n=4 animals; Uninjured+Ibuprofen: n=4; Uninjured+Ibu+Carnitine: n=3; SBI: n=3; SBI+Ibuprofen: n=3; SBI+Ibu+Carnitine: n=3. Ibuprofen and carnitine were supplemented via drinking water at 0.6mg/mL and 1mg/mL respectively. Due to poor solubility of ibuprofen in water, ibuprofen was converted to sodium ibuprofenate prior to addition to drinking water. Briefly, ibuprofen (1 equiv.) was dissolved in a minimal volume of ethanol and aqueous 1M NaOH solution (1 equiv.) was added dropwise at room temperature while stirring. After 15–30 minutes of stirring, sodium ibuprofenate formed, and the solvent was then removed under reduced pressure using a rotary evaporator. The solid ibuprofen sodium salt was then dissolved in drinking water to a final concentration of 0.6 mg/mL. All injured groups drank equivalent amounts of water (**Supplementary Table 5**).

Fourteen days after delivery, the pubocaudalis (PCA) muscle was harvested for histologic analysis. Immediately after dissection, the muscle was embedded in Optimal Cutting Temperature (OCT) compound and snap-frozen using liquid nitrogen-cooled isopentane (2-methylbutane), then stored at −80°C until sectioning. 10 µm cross-sectional sections were obtained using a cryostat. Sections were mounted on positively charged glass slides and air-dried briefly at room temperature before being returned to −80°C storage until staining.

Antibody staining was performed using a three day protocol. Frozen slides were equilibrated to room temperature for 15 minutes, then. hydrophobic barriers were drawn around each tissue section using an ImmunoEdge pen. Slides were briefly dipped once in distilled water and fixed by gentle immersion in 10% formalin for 15 minutes at room temperature. Following fixation, slides were rinsed three times in 1× phosphate-buffered saline (PBS) for 2 minutes each. Tissue permeabilization was performed by incubating slides in 0.2% Triton X-100 in PBS for 10 minutes at room temperature, followed by three 1-minute rinses in PBS. Next, sections were incubated overnight at 4°C with FabFragment reagent diluted 1:100 in 1× PBS and incubated in a blocking buffer consisting of 10% goat serum and 3% bovine serum albumin (BSA) in PBS for 1 hour at room temperature. The primary antibody used was Laminin Polyclonal Antibody (Thermofisher, catalog #: PA1-16730). It was diluted 1:500 in blocking buffer and applied to each section at approximately 200 µL per slide, ensuring complete tissue coverage. Slides were incubated overnight at 4°C, then gently washed in 1× PBS, including a 10-minute immersion step. The secondary antibody (goat anti-rabbit Alexa Fluor 488, Thermofisher, catalog #: A11008) was diluted 1:200 in blocking buffer and applied to sections for 1 hour at room temperature, protected from light. Following secondary antibody incubation, slides were washed by five dips in 1× PBS followed by a 10-minute immersion in PBS, then immersed in Milli-Q water for 10 minutes. Slides were blotted dry with lint-free tissue, and coverslips were mounted using Vectashield mounting medium with DAPI nuclear stain.

Muscle fiber cross-sectional area (CSA) was quantified from immunofluorescence images of Laminin-stained muscle cross-sections using QuPath (v0.7.0) and the QuPath-Cellpose extension (v0.12.0, QuPath-BIOP), following an approach adapted from Reinbigler et al. (2022)^61^. Individual muscle fibers were segmented using the Cellpose deep learning model (cyto3) applied to the green fluorescence channel. Fibers intersecting the image boundary were automatically excluded to prevent artificially reduced CSA measurements from truncated fibers. Shape measurements, including area, perimeter, and circularity, were computed for all detected fiber annotations. The QuPath pipeline is available at https://github.com/hannahheathomics/multiplex_synthesis.

Centrally nucleated fiber analysis was performed using CellProfiler (version 4.2.8), following the pipeline described by Lau et al. (2018)^62^ with modifications. Briefly, muscle fibers were segmented from the inverted Laminin channel using intensity-based thresholding (manual threshold: 0.5, three-class) with intensity-based declumping and local maxima suppression (minimum distance: 15 pixels). Nuclei were segmented from the DAPI channel and shrunk to single points before being assigned to their parent fiber. A fiber was classified as centrally nucleated if it contained at least one nucleus. Centrally nucleated fiber rate was calculated as the percentage of fibers classified as centrally nucleated per region of interest. The CellProfiler pipeline is available at https://github.com/hannahheathomics/multiplex_synthesis.

## Analysis of public GNPS/MassIVE datasets

The multiplex synthetic library created was used to analyze multiple datasets: (1) inflammatory bowel disease ([MSV000082094](https://massive.ucsd.edu/ProteoSAFe/dataset.jsp?task=b68a5f90cb4f4743978ba2dc0600f7f2), fecal samples); (2) a pediatric IBD cohort ([MSV000097610](https://massive.ucsd.edu/ProteoSAFe/dataset.jsp?task=65cc240951884d4dadbfe2edf4172e03), fecal samples); (3) a rheumatoid arthritis cohort ([MSV000084556](https://massive.ucsd.edu/ProteoSAFe/dataset.jsp?task=96b37796c1214cbb9151d2bd676b31c1), fecal samples). Each public dataset was downloaded and processed using MZmine using the batch workflow for feature finding and detection. An example of a.mzbatch file containing detailed parameter settings can be found in https://github.com/VCLamoureux/synthesis_multiplex. The output files generated using the batch processing workflow (a csv file with peak areas and an mgf file with MS/MS information associated with each feature) were used as input for feature-based molecular networking (FBMN) in GNPS2 and ran against the multiplex synthetic library. The FBMN parameters were set for each dataset with a cosine of 0.7, precursor and fragment ion tolerance of 0.02 Da, filters set to off, and a number of matched peaks for networking and library search, set to 4. For each dataset, the quantification table (generated via MZmine), the annotation table (FBMN workflow), and the metadata was imported in RStudio for data formatting. The formatted data tables were exported into a csv file for boxplots creation using Python scripts (see Code availability).

## Computational infrastructure

This is an expanded description of the computational infrastructure that was developed to enable this project. MASST^7^ (Mass Spectrometry Search Tool) queries now run on a virtual machine equipped with two 64-core AMD Milan EPYC 7713 processors and 2 TB of RAM, with public metabolomics data indices hosted on four NVMe Solidigm D5-P5336 SSDs configured in a RAID ZFS striped array. We refer to these as fast MASST or FASST^63^ queries. The GNPS2 platform has expanded to operate across five virtual machine servers: two with dual 64-core AMD Milan EPYC 7713 processors and 2 TB of RAM, and three with dual 16-core Intel E5-2683 v4 CPUs and 768 GB of RAM. Storage is provided by two arrays: one comprising 24 × 7.68 TB SATA SSDs (184 TB) and another with 8 × 30 TB NVMe SSDs (240 TB). All servers are interconnected via a 10 Gbit network.

## Chemical synthesis

NMR spectra were collected at 298 K on a 600 MHz Bruker Avance III spectrometer fitted with a 1.7 mm triple resonance cryoprobe with z-axis gradients. (^1^H NMR: MeOD (3.31), CDCl_3_ (7.26) at 600 MHz. 5-ASA-phenylpropionic acid spectra was taken in MeOD with shifts reported in parts per million (ppm) referenced to the proton of the solvent (3.31), and Ibuprofen-carnitine spectra was taken in CDCl_3_ with shifts reported in parts per million (ppm) referenced to the proton of the solvent (7.26), Coupling constants are reported in Hertz (Hz). Data for ^1^H-NMR are reported as follows: chemical shift (ppm, reference to protium; s = single, d = doublet, t = triplet, q = quartet, dd = doublet of doublets, m = multiplet, coupling constant (Hz), and integration).

## Multiplex reactions

The synthesis procedures for 5-ASA-phenylpropionic acid and ibuprofen-carnitine are detailed below. Additionally, the complete set of multiplex library reaction protocols, including all reagents, and conditions are provided in **Supplementary information**.

## 5-ASA-phenylpropionic acid

Solid 5-aminosalicylic acid (2 mmol, 100 mg, 1 eq.) and 3 mL of THF were added sequentially to a 20 mL glass vial with a stir bar and the reaction was placed in an ice/water bath, neat triethylamine (Et_3_N) (2.41 mmol, 419 µL, 1.2 eq.) and phenylpropionyl chloride (2 mmol, 307 μL, 1 eq.) were added under inert atmosphere, and the solution was stirred for 5 h at 23 °C. The mixture was concentrated using a rotary evaporator and the crude material was purified using a CombiFlash NextGen 300+ with reversed phase column C18 15.5 g Gold at a flow rate 13 mL per min with H_2_O (Solvent A) and ACN (solvent B) with the following gradient: 0-5 min, 5% B; 5-14 min, 40% B; 14-20 min 40% B; 20-25 min, 80% B. 5-ASA-phenylpropionic acid eluted at 15 min, 40% B. ^1^H-NMR (MeOD) δ 2.61 (t, 3H), 9.98 (t, 3H), 6.76 (2, 1H), 7.11-7.30 (m, 5H), 7.41 (d, 1H), 7.99 (d, 1H) (^1^H-NMR spectra is available 10.5281/zenodo.17519052).

## Ibuprofen-carnitine

Solid ibuprofen (4.85 mmol, 1 g, 1 eq.) and 3 mL of THF were added to a 20 mL vial with a stir bar and the reaction was placed in an ice/water bath, neat ethyl-chloroformate (5.82 mmol, 559 µL, 1.2 eq.) and triethylamine (7.27 mmol, 1.01ml, 1.5 eq.) were subsequently added, and the solution was stirred for 1.5 h, solid carnitine (4.85 mmol, 958 mg, 1 eq.) dissolved in 2 mL THF was subsequently added to the ibuprofen mixture. The reaction was removed from the ice/water bath and stirred overnight at 23 °C. The mixture was concentrated en vaccuo and purified using a CombiFlash NextGen 300+ with reversed phase column C18 15.5 g Gold at a flow rate13 mL per min with H_2_O (Solvent A) and ACN (solvent B) using the gradient: 0-2 min, 5% B; 3-10 min, 20-40% B; 11-14 min 60% B; 15-17 min, 80% B. Ibuprofen-carnitine eluted at 3-10 min, 20% B. ^1^H-NMR (CDCl3) δ 0.86 (m, 6H), 1.73 (m, 3H), 1.18 (m, 1H), 2.22 (m, 2H), 2.42 (dd, 2H), 2.85-3.11 (m, 9H), 3.42 (m, 1H), 5.5 (m, 1H), 7.08-7.14 (m, 4H) (^1^H-NMR spectra is available 10.5281/zenodo.17519052).

## Quantification

Quantification of Ibuprofen-carnitine and 5-ASA-phenylpropionic acid was performed from urine sample and fecal sample, respectively.

The LC-MS/MS method used for the analyses of the method validation and quantification was the same as previously described in LC-MS/MS data collection. The analytical method was performed according to the International Conference on Harmonization (ICH) guidelines^64^ for ibuprofen-carnitine and 5-ASA-phenylpropionic acid. The method was validated based on the evaluation of the following parameters: specificity, precision (repeatability and intermediate precision), linearity, limit of detection (LOD), limit of quantification (LOQ), and accuracy. A matrix match calibration curve was created by spiking pool urine (ibuprofen-carnitine) and pool fecal (5-ASA-phenylpropionic acid) into calibrates to create a matrix match calibration curve for quantitation. Detailed information regarding the methodology used for each of them is described below. The validation was performed using Rise Plus Urobiome samples of human urine MSV000096359 that would contain the ibuprofen-carnitine compound and Crohn’s cohort MSV000099375 that contains the 5-ASA-phenylpropionic acid. Peak area for ibuprofen-carnitine and 5-ASA-phenylpropionic acid was extracted using Skyline^65^ (version 23.1). The method employed reached the acceptance criteria specified for each parameter of ibuprofen-carnitine (**Supplementary Table 6**), and 5-ASA-phenylpropionic acid (**Supplementary Table 7**). For quantification in biological samples, one sample of the Crohn’s cohort and one sample of the Rise Plus Urobiome study with the highest peak area was injected in the validated method (samples were resuspended in 100 μL of 50/50 MeOH/H_2_O containing 1 μM of sulfamethazine). For the calculation of the amounts in the samples, it was estimated that 200 uL of urine sample and 54 mg fecal samples would be the starting material, and the extraction yield was also extrapolated to 100%.

## Specificity

The specificity was determined by injecting a blank solution containing only the internal standard (sulfadimethazine), and an injection of a solution containing all the ibuprofen-carnitine (n=3). The relative standard deviation (RSD) was calculated based on each peak’s retention time in the Rise Plus Urobiome and fecal samples. The MS and MS/MS spectra confirmed the specificity and identity of these compounds. The retention times of the peak of interest were as follows: Ibuprofen-carnitine at 2.09 min and 5-ASA-phenylpropionic acid at 4.52 min. These compounds didn’t show interferences compared to the solution containing only the mixture of standards.

## Linearity

The linearity of the method was determined by calibration curves in concentration ranges comprising each compound at the samples of interest. A stock solution containing 1mM of each Ibuprofen-carnitine and 5-ASA-phenylpropionic acid was prepared in 50/50 MeOH/H_2_O, followed by serial dilutions and used to acquire calibration curves for all the compounds simultaneously. From this solution, 7 points were prepared with levels ranging from 10 nM to 1 μM for Ibuprofen-carnitine and 100 nM to 2 μM for 5-ASA-phenylpropionic acid with each spike with urine matrix Ibuprofen carnitine and fecal matrix for 5-ASA-phenylpropionic acid. Each concentration level was injected in triplicates and the analytical curves were built based on the nominal concentrations, and the average between the ratios of each compound and the internal standard used (Ratio = A_compound_/A_IS_). A polynomial equation was obtained for each curve, and the correlation coefficients were calculated for each compound.

## Limit of detection and limit of quantification

LODs and LOQs were estimated by the mean of the slopes (S) and the standard deviation of the y-intercept (y). These limits were calculated by the following equations: LOD = (3.3∗y)/S and LOQ = (10∗y)/S. All the slopes, intercepts, LODs, and LOQs are shown in **Supplementary Tables 6-7**.

## Accuracy and Precision

The accuracy and precision of the method was determined by recovery analyses. For this, known amounts of the solution containing the standards were spiked to the sample P1-D-6_2_5753 for 5-ASA-phenylpropionic acid and sample STD_SPK_urine for Ibuprofen-carnitine solutions in two different concentrations (low and high) considering the predetermined calibration curve and concentration range. Three replicates for each level were injected and analyzed in the validated method. The accuracy was determined by the difference between the theoretical and experimental concentration values and the values were within the acceptance range of 80–120% and the precision by coefficient variation (CV).

## Data availability

All data in this study are publicly available and accessible on GNPS/MassIVE. The multiplex synthesis library is available at https://external.gnps2.org/gnpslibrary (“MULTIPLEX-SYNTHESIS-LIBRARY”, 6 partitions in total). All untargeted metabolomics LC-MS/MS data have been deposited to GNPS/MassIVE under the accession numbers MSV000097885, MSV000097874, MSV000097869, MSV000094559, MSV000094447, MSV000094393, MSV000094391, MSV000094382, MSV000094337, MSV000094300, MSV000098637 (bile acid), MSV000098628 (small molecules), MSV000098639 (drug compounds), MSV000098640 (peptides), MSV000096359 (Rise Plus Urobiome samples), and MSV000099150 (urine from 9 pregnant women), MSV000099374 (data files for standards and biological samples for retention time matching of 5-ASA, ibuprofen and succinic acid conjugates), MSV000099375 (5-ASA quantification data), MSV000099556 (ibuprofen quantification data), quantification datasets for OCTN2 and Ibuprofen carnitine (MSV000101313). The job link for searching the multiplex synthesis library against existing GNPS libraries is available at https://gnps2.org/status?task=0e77aa138fc2473ab8a801a8d59905e6. The classical molecular network of synthetic MS/MS spectra that are exclusively present in humans is available at https://gnps2.org/status?task=a6b9129f880146b0aef3168855c32713. The FBMN jobs of three datasets used for 5-ASA-phenylpropionic acid can be accessed using the following links: IBD dataset (MSV000082094): https://gnps2.org/status?task=5f230f976ccb4f19aa94d59407468138; Pediatric IBD cohort (MSV000097610): https://gnps2.org/status?task=023988a1842146d6a5b2ba87a3212598; Rheumatoid arthritis cohort (MSV000084556): https://gnps2.org/status?task=16da6e571d574a829e3de75dd610bc97. Histology image files are available at https://doi.org/10.5281/zenodo.19637023.

## Code availability

Source codes for all the data analyses applied on the multiplex synthesis library can be found at https://github.com/Philipbear/multiplex_synthesis. The tool for generating SMILES strings of multiplex synthesis products can be accessed at https://autosmiles.streamlit.app/rxnSMILES. All scripts used for quantification are available at https://github.com/abubakerpatan/Quantification-Script. The codebase of the MS/MS library generation workflow is available at https://github.com/Wang-Bioinformatics-Lab/Reverse_metabolomics_library_generation. All scripts used to generate the figures in this study can be accessed from GitHub (https://github.com/Philipbear/multiplex_synthesis and https://github.com/VCLamoureux/synthesis_multiplex).

## Use of AI and Software with AI for the research

In the Dorrestein Lab, the use of AI and AI-enabled tools, including generative AI, large language models (LLMs), and software that leverages these technologies, both free and commercial, is encouraged across all aspects of research. These tools support a wide range of activities, including literature searches, data analysis, scripting, coding, text editing, and figure concept development. All figures and analyses are original. To ensure transparency and reproducibility, all raw data, derived data tables and code used in this research are made accessible and linked with this manuscript.

## Acknowledgements

PCD acknowledges NIDDK R01DK136117, U24DK133658, and EnvedaGives Scientific Research Fund and their support for the Human Chenome Project for enabling this work. LAB is supported by the Reproductive Scientist Development Program (NIH K12HD000849) and the FIRST Program (NIH U54CA272220). MW acknowledges NIGMS U24DK133658. AP was supported by R01DK136117. SX was supported by BBSRC/NSF award 2152526. VCL was supported by Fonds de recherche du Québec – Santé (FRQS) postdoctoral fellowship (335368) and from Natural Sciences and Engineering Research Council of Canada (NSERC) postdoctoral fellowship (598938). NB and JC were partially supported by NIDDK U24DK133658 and NIGMS R24GM148372. HM-R is supported by U24DK133658. LCD is supported by the DDRA (Doctoral Dissertation Research Award) from the Fulbright U.S. Student Program, which is sponsored by the U.S. Department of State and the Fulbright Brazil Commission. AMC-R and PCD were supported by the Gordon and Betty Moore Foundation grant GBMF12120 and https://doi.org/10.37807/GBMF12120. Reproductive Scientist Development Program Grant supports LAB, ACTRI Pilot Project Grant to LAB. KMG acknowledges support from the NIH: UC2HD113474. HG acknowledges support from Crohn’s and Colitis Foundation of America (CCFA, Grant ID:1243263) and Helmsley Foundation and from Eric & Wendy Schmidt AI in Science Postdoctoral Fellowship. AJ acknowledges financial support from the Howard Hughes Medical Institute through the Hanna H. Gray Fellowship. The NMR data reported in this publication was supported by the Office of the Director of the National Institutes of Health under award number S10 OD032266. The content is solely the responsibility of the authors and does not necessarily represent the official views of the National Institutes of Health. YEA acknowledges funding through the Austrian Academy of Sciences (ÖAW) through APART-USA.

## Author contributions

PCD conceptualized the idea. LAB conceptualized the muscle injury study and obtained urine samples. AP led the synthesis part of the project. SX developed the codes for library generation. SX and VCL performed data analysis. AP, SX and JZ created the MS/MS libraries. JY, ED and KMG developed the OCTN2 assay, YEA, MW developed and managed indexing, metadata harmonization and enabled fast MASST in GNPS2. JZ, LA, AM, MAP, ACF, EO and HH performed mice work and histology. RAM and AJ developed the Boltz-2 ensemble co-folding predictions model. JC, JD, NB enabled the hardware and software for MassIVE in USIs in GNPS. AP, VCL, JA, WG and IM performed the LC-MS/MS data collection. AP, ZH, VD, DL, TB, SG, NW, VN, and WT worked on synthesis. PR developed the SMILES generation tool. CL, AP worked on the SMILES generation. ZH, VD, AA, and LCD helped with sample preparation. HG, WDGN, HNZ, SZ, KEK, HM-R, AMCR tested synthesis libraries. JY, ED and KMG performed transporter assays. AP, SX, VCL, and PCD drafted the manuscript. DS, LAB and PCD supervised the project. All authors reviewed and approved the manuscript.

