## Supplementary information for "Charting the undiscovered metabolome with synthetic multiplexing implicates ibuprofen-carnitine in myotoxicity"

**Affiliations**:
[1]Skaggs School of Pharmacy and Pharmaceutical Sciences, University of California San Diego, La Jolla, CA, USA
[2]Department of Medicine, University of California, San Diego, San Diego, CA
[3]Department of Pharmacology, University of California San Diego, La Jolla, CA, 92093, USA
[4]Collaborative Mass Spectrometry Innovation Center, Skaggs School of Pharmacy and Pharmaceutical Sciences, University of California San Diego, La Jolla, CA, USA
[5]Center for Microbiome Innovation, University of California San Diego, La Jolla, CA, 92093, USA
[6]Department of Obstetrics, Gynecology, and Reproductive Sciences, Division of Urogynecology and Reconstructive Pelvic Surgery, University of California San Diego, La Jolla, CA 92093, USA
[7]Department of Computer Science, University of California Riverside, Riverside, CA, USA
[8]Department of Computer Science and Engineering, University of California at San Diego, La Jolla, CA 92093, USA
[9]Center for Computational Mass Spectrometry, University of California San Diego
[10]Laboratório de Métodos de Extração e Separação (LAMES), Instituto de Química (IQ), Universidade Federal de Goiás (UFG), Campus II – Samambaia, Goiânia, GO, 74045-155, Brasil
[11]Department of Chemistry and Biochemistry, UC San Diego, La Jolla, California, USA
[12]Department of Bioengineering and Therapeutic Sciences, University of California, San Francisco, USA.
[13]Department of Orthopedic Surgery.
[14]Amazon Integrated Metabolomics Center (CIMAZON), Institute of Natural and Exact Sciences, Federal University of Pará, Rua Augusto Corrêa, 01 - Guamá 66075-110, Belém, PA, Brazil
[15]BOKU University, Department of Chemistry, Institute of Analytical Chemistry, Muthgasse 18, 1190, Vienna, Austria

[†] Abubaker Patan, Shipei Xing, and Vincent Charron-Lamoureux contributed equally to this work.

**Disclosures:** PCD is an advisor and holds equity in Cybele, BileOmix, Sirenas and a scientific co-founder, advisor, holds equity and/or received income from Ometa, Enveda, and Arome with prior approval by UC San Diego. PCD also consulted for DSM animal health in 2023. MW is a co-founder of Ometa Labs.

**Table of Contents**

**Supplementary Table 1**. It lists all reactants used in the multiplex synthesis experiments. Each row corresponds to a distinct compound and includes detailed chemical and structural information. The table provides the following columns: compound_name (the name of the derived or related reactant), SMILES (the original Simplified Molecular Input Line Entry System representation), corrected_SMILES (curated SMILES strings adjusted for syntax and structure accuracy), formula (the molecular formula), neutralized_formula (the formula standardized to a neutral charge state), exact_mass (monoisotopic mass), np_class and np_superclass (natural product classification levels), np_pathway (the biosynthetic or chemical pathway classification), inchikey (a standardized InChIKey for compound identification), and 2d_inchikey (an InChIKey truncated to represent 2D connectivity only). This table serves as a comprehensive reference for all chemical entities involved in the multiplex synthetic reactions performed in MS\MS spectral library creation.

**Supplementary Table 2.** It provides a comprehensive list of the 4,000 chemical reactions conducted as part of this study. Each reaction is uniquely identified by a Reaction_ID and described by its Reaction_name, with additional details on the sequential synthetic steps (Step_2, Step_3, and Step_4). The table includes direct access to experimental records via the Signal_notebook_link (which can be only accessed by UCSD user credentials) where all reactions are documented; an example of this document is 5_ASA_fatty-acid.pdf. Analytical information is provided through the Run_on_MS (mass spectrometry) and the corresponding MassIVE_ID for raw data access. The Processing_software column indicates the tools used for library creation. The table also tracks spectra curation into libraries, indicating whether a Library_Created, This table serves as a detailed experimental and analytical record supporting reproducibility and data transparency.

**Supplementary Table 3.** It contains the results of the overlap analysis between the multiplexed synthetic MS/MS library generated in this study and several major public structural databases, including PubChem, HMDB, GNPS, PubChemLite, NORMAN, and FooDB.

**Supplementary Table 4.** It contains the 2,679 spectral matches of the multiplexed synthetic library that exclusively matched to human data.

**Supplementary Table 5.** It contains water consumption by mice that contained Ibuprofen and carnitine that were supplemented via drinking water.

**Supplementary Table 6.** It contains the Quantification, Specificity, Linearity, Limit of detection and limit of quantification, Accuracy and Precision data for Ibuprofen-carnitine

**Supplementary Table 7.** It contains the Quantification, Specificity, Linearity, Limit of detection and limit of quantification, Accuracy and Precision data for 5-ASA-phenylpropionic acid.

**Supplementary Table 8.** Carnitine and Ibuprofen-carnitine conjugate quantification in carnitine and ibuprofen exposed mice, injured and uninjured.

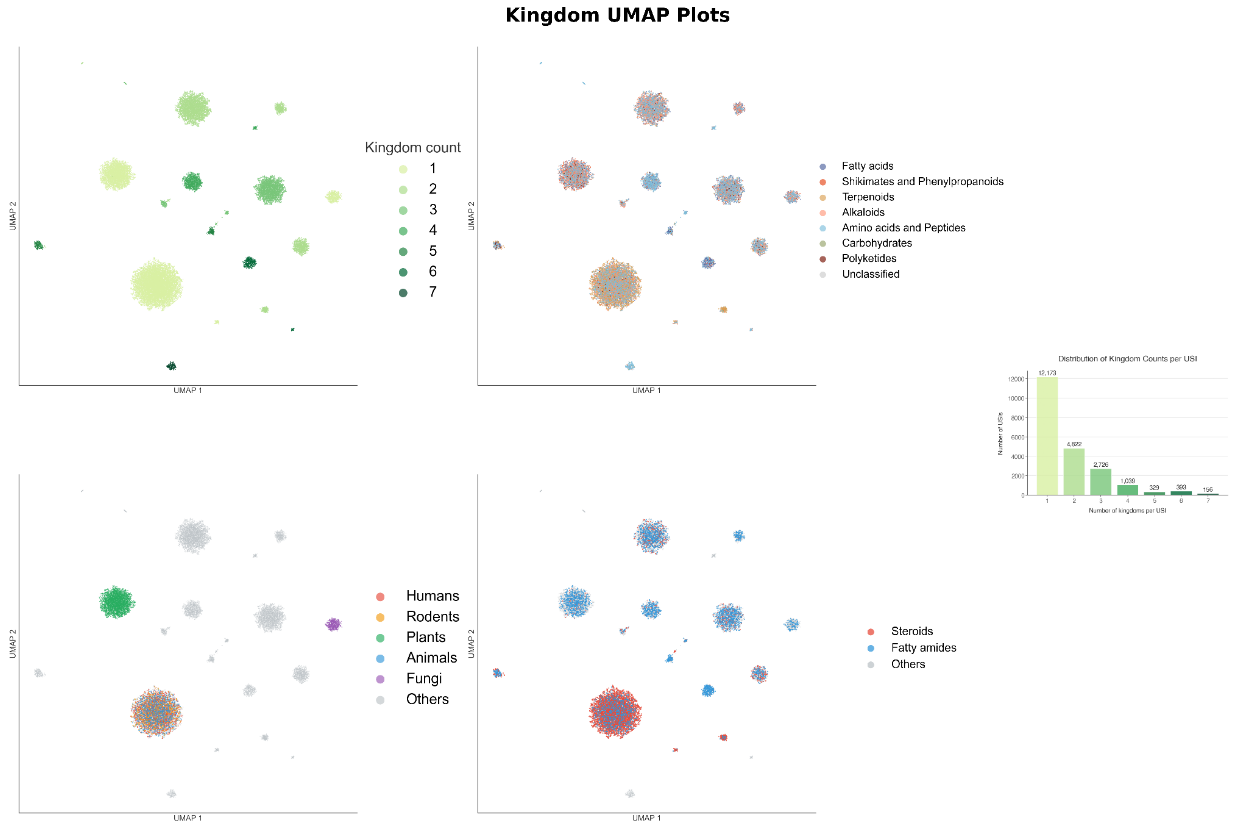


**Supplementary Figure 1 | UMAP visualizations of the kingdom distribution of matched spectra across public datasets.**

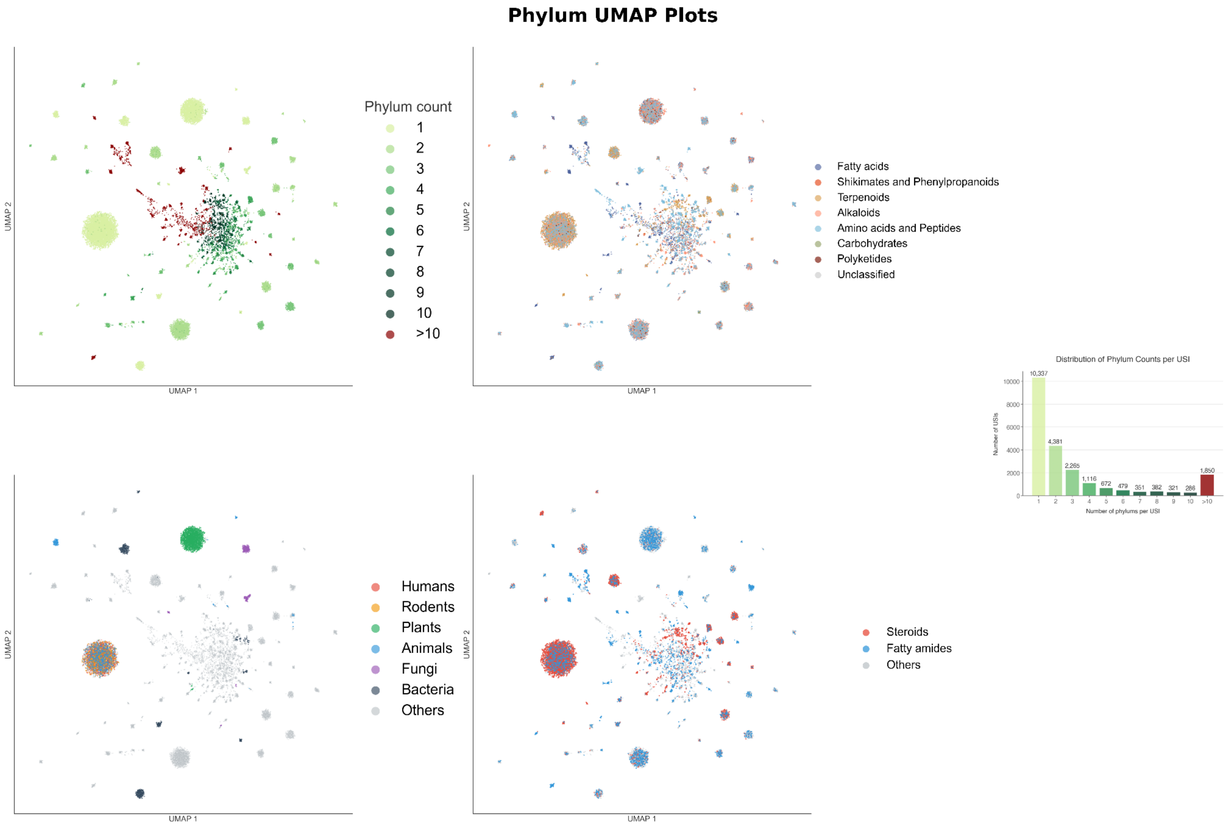


**Supplementary Figure 2 | UMAP visualizations of the phylum distribution of matched spectra across public datasets.**

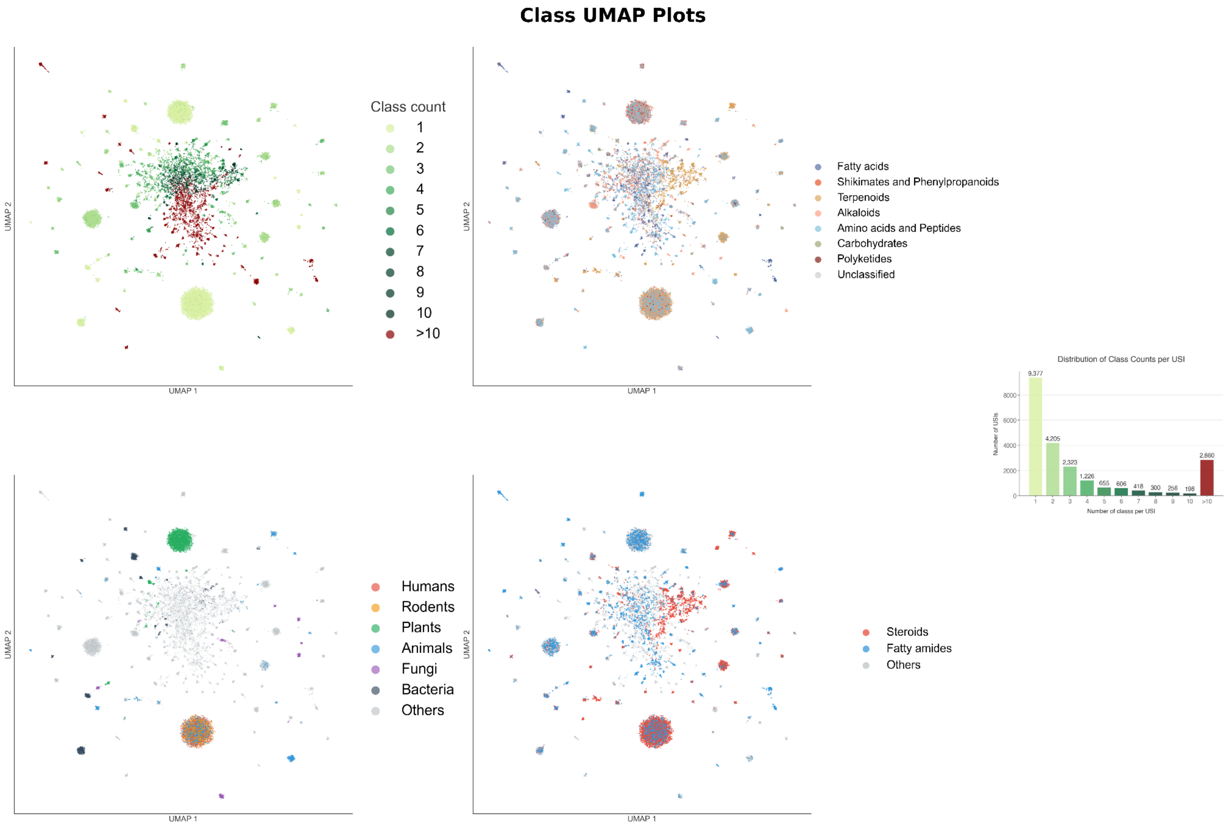


**Supplementary Figure 3 | UMAP visualizations of the class distribution of matched spectra across public datasets.**

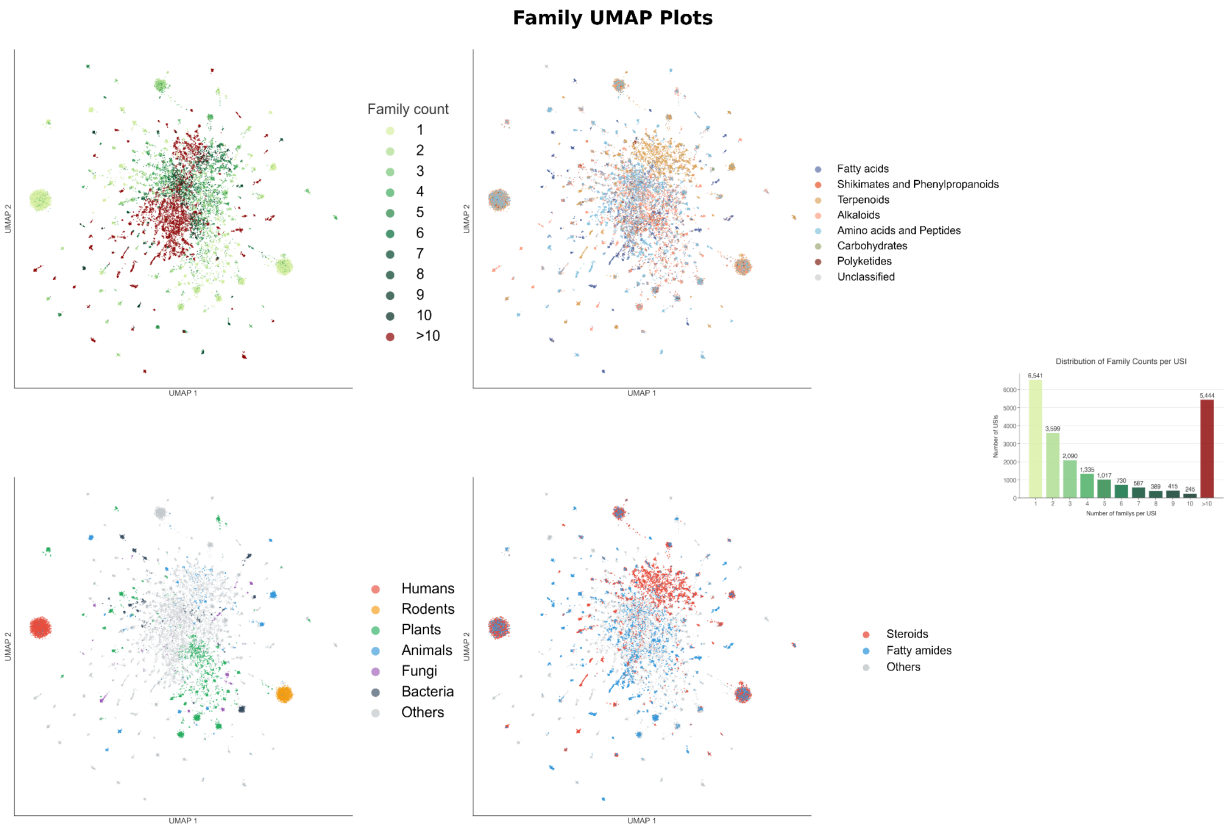


**Supplementary Figure 4 ┃ UMAP visualizations of the family distribution of matched spectra across public datasets.**

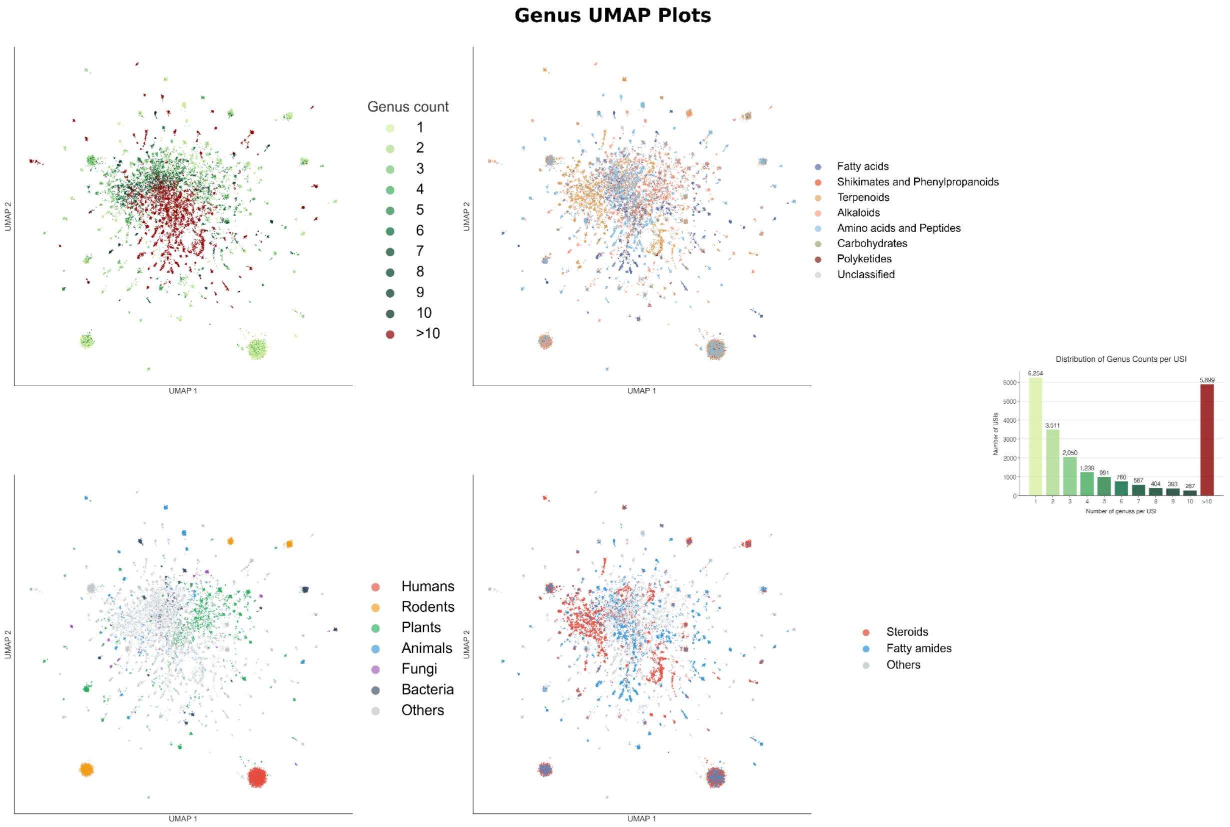


**Supplementary Figure 5 | UMAP visualizations of the genus distribution of matched spectra across public datasets.**

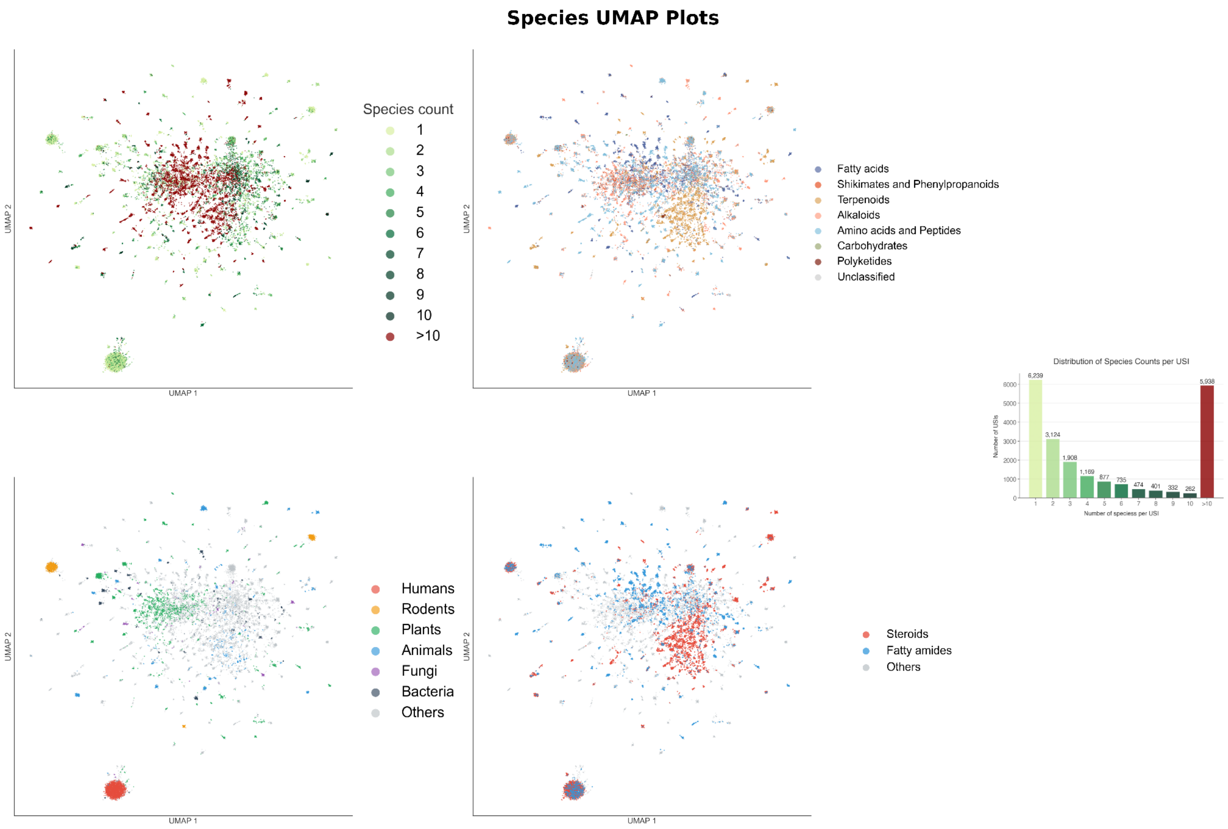


**Supplementary Figure 6 | UMAP visualizations of the species distribution of matched spectra across public datasets.**

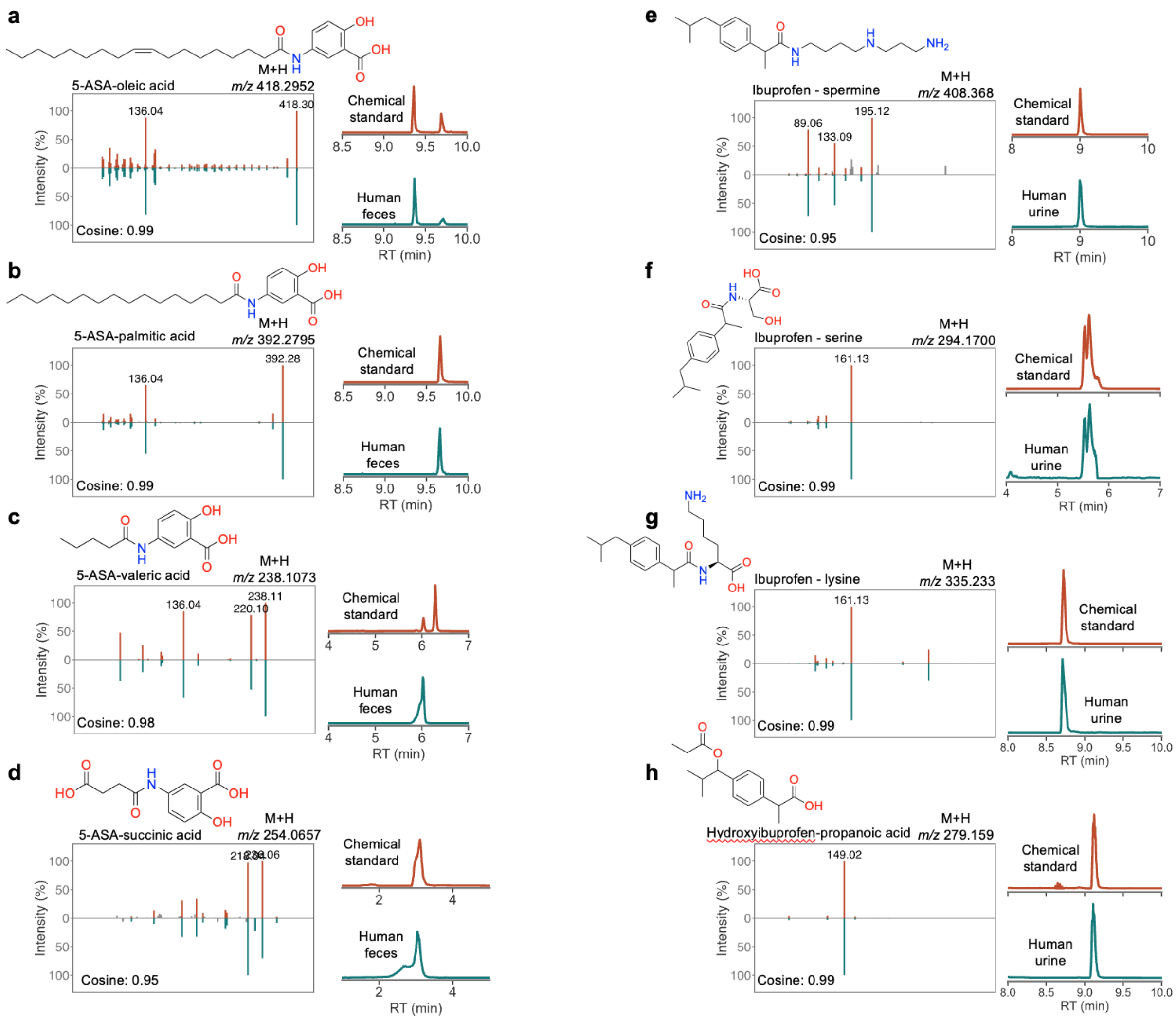


**Supplementary Figure 7 | Structure validation of 5-ASA and ibuprofen drug metabolites.** MS/MS spectral matching and retention time alignment of **a)** 5-ASA-oleic acid, (should be interpreted as a 5-ASA-C18:1 fatty acid conjugate as other isomers are possible) **b)** 5-ASA-palmitic acid (C16:0 fatty acid), **c)** 5-ASA-valeric acid, (C5:0 fatty acid) **d)** 5-ASA-succinic acid, **e)** ibuprofen-spermidine, **f)** ibuprofen-serine, **g)** ibuprofen-lysine, and **h)** hydroxyibuprofen-propanoic acid.

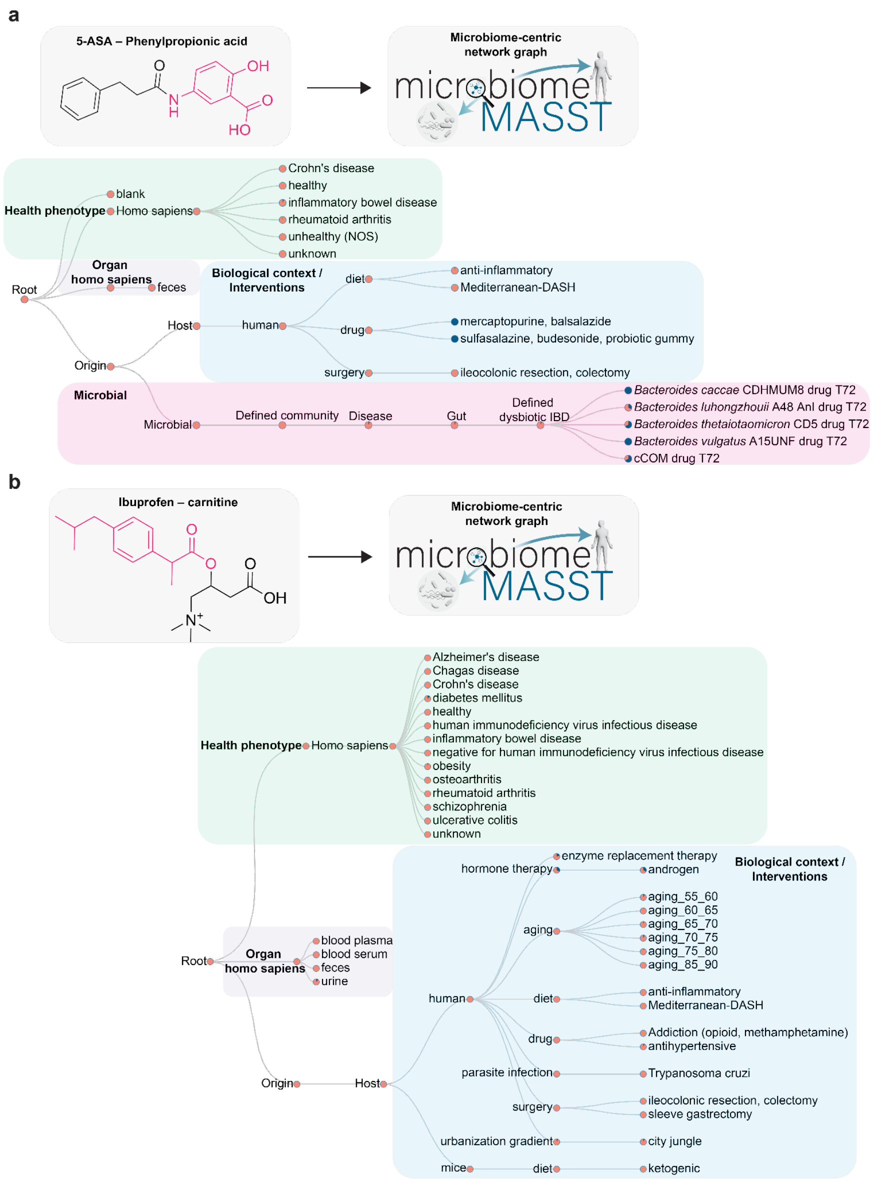


**Supplementary Figure 8 ┃ The distribution of 5-ASA-phenylpropionic acid and ibuprofen-carnitine in host biology. a)** The universal spectrum identifier (USI) of 5-ASA-phenylpropionate retrieved from the GNPS2 library was used as input in microbiomeMASST. MicrobiomeMASST results showed that 5-ASA-phenylpropionate was detected in bacterial culture supplemented with the substrates 5-ASA and phenylpropionate, exclusively observed in humans under diets, drugs, surgery interventions, and in health conditions including Crohn's disease, inflammatory bowel disease, and rheumatoid arthritis. **b)** Ibuprofen-carnitine was detected in several health conditions including obesity, osteoarthritis, and ulcerative colitis, across several biological contexts such as aging and parasite infection, and found in blood, feces, and urine.

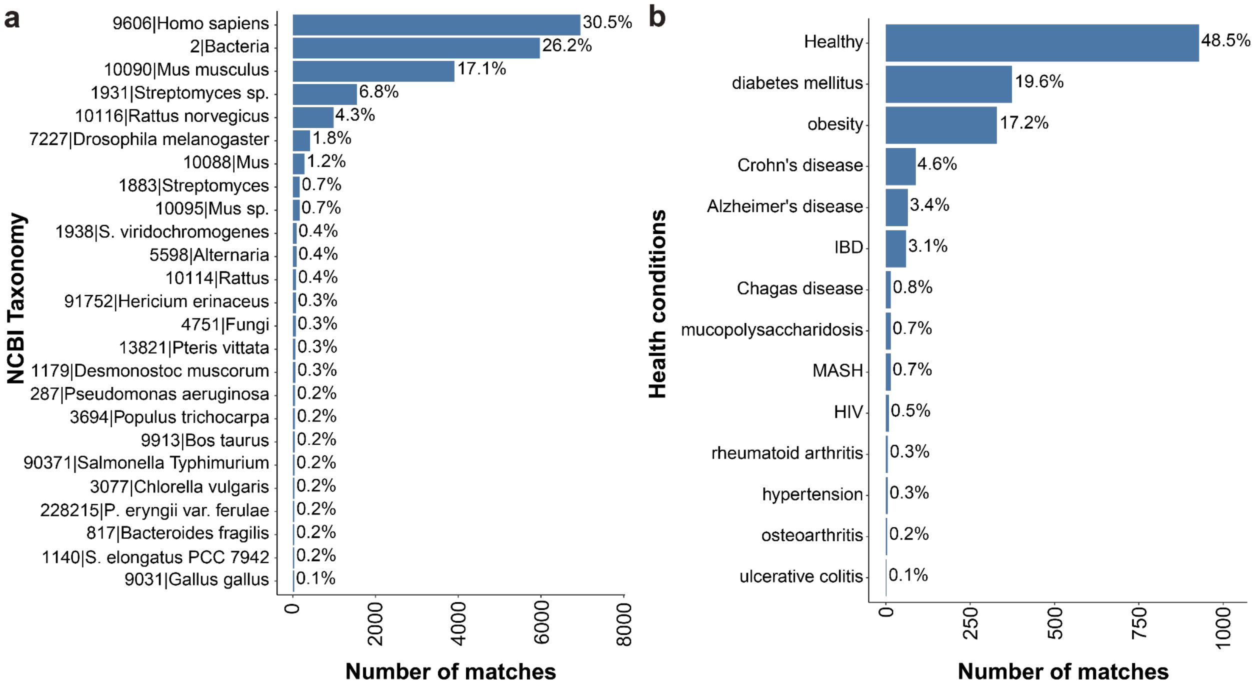


**Supplementary Figure 9 | Prevalence of succinic acid conjugates across taxonomy and health conditions. a)** Distribution of succinic acid conjugates matched across NCBI Taxonomy using the Pan-ReDU database. The bars show the number of matches. On the right side of the bars, the percentage of all matches is shown. **b)** Distribution of succinic acid conjugates matched across health conditions. The bars show the number of matches and the right labels show the percentage of only human matches. IBD, inflammatory bowel disease; MASH, metabolic dysfunction-associated steatohepatitis; HIV, human immunodeficiency virus.

**Multiplex synthesis**

Multiplex syntheses were conducted utilizing standard chemical methodologies. The reaction parameters, such as temperature, reagent concentration, solvent composition, and reaction duration, were systematically refined to ensure consistent reaction efficiency across all targets. Subsequent adjustments to the method were implemented based on the solubility of substrates, their chemical reactivity, and sensitivity. The polarity of solvents and buffer systems were modified to enhance substrate dissolution and stability, while the concentrations of catalysts were optimized to improve reaction yields. These adjustments guaranteed that the multiplex synthesis was carried out efficiently, achieving high yield and selectivity.

**Amidation (a).** A carboxylic acid containing compound (1 eq.) and 2 mL of DMF were added to a 20 mL scintillation vial with a magnetic stir bar. To this solution, solid EDC (1.2 eq.) and neat DIPEA (1.5 eq.) were subsequently added, and the solution was stirred at 23°C. After 15 minutes, a mixture containing each amine (0.01 eq.), DMAP (0.2 eq.) were added, and the reaction was stirred for 14 hours. The mixture was then concentrated *in vacuo* and an LC-MS/MS sample was prepared.

**Amidation (b).** A carboxylic acid containing compound (1 eq.) and 2.5 mL of anhydrous THF were added to a 20 mL scintillation vial with a magnetic stir bar The solution was cooled to 0°C with an ice bath. To this solution, ethyl chloroformate (1.2 eq.) and $Et_3N$ (1.2 eq.) were added, and the solution was stirred at 0°C for 1.5 hours. A mixture containing 0.01 eq of each amine was prepared in 2.5 mL of water with 1.5 eq of NaOH. After 1.5 hours of stirring, the amine mixture was added to the activated carboxylic acid mixture. The reaction was then stirred for 3 h and allowed to gradually warm to 23°C, yielding a clear, homogeneous solution. The mixture was then concentrated *in vacuo* and an LC-MS/MS sample was prepared.

**Amidation (c).** An amine containing compound (1 eq.) and 2.5 mL of anhydrous DCM were added to a 20 mL scintillation vial with a magnetic stir bar The solution was cooled to 0°C with an ice bath. To the cooled solution, $Et_3N$ or pyridine (1.2 eq.) were subsequently added, and a mixture of acyl chlorides (0.01 eq. per compound) was added dropwise to the mixture. The mixture was stirred for 2 to 3 hours and allowed to gradually warm to 23 °C. The mixture was then concentrated *in vacuo,* and an LC-MS/MS sample was then prepared from the concentrated mixture.

**Amidation (d).** A carboxyl compound (1 eq.) was dissolved in 2.5 mL of DMF in a scintillation vial with a magnetic stir bar. To the solution, (10 eq.) of ghosez's reagent was  added.This was then stirred until the reaction mixture turned yellow, then (2 eq.) of neat pyridine and the amine containing compounds were added. The reaction mixture was then stirred  2-3 hr. The mixture was then concentrated *in vacuo* and an LC-MS/MS sample was prepared.

**Esterification (a).** A carboxylic acid compound (1 eq.) and 2 mL of DMF were added to a 20 mL scintillation vial with a magnetic stir bar. To this solution, solid EDC (1.2 eq.) was subsequently added, and the solution was stirred at 23°C. After 15 minutes, A mixture containing 0.01 eq of each alcohol , and DMAP (0.5 eq.) were added, and the reaction was stirred for 14 hours. The mixture was then concentrated *in vacuo* and an LC-MS/MS sample was prepared.

**Esterification (b).** An alcohol compound (1 eq.) and 2.5 mL of anhydrous DCM were added to a 20 mL scintillation vial with a magnetic stir bar. The solution was cooled to 0°C with an ice bath. To this solution, either $Et_3N$ or pyridine (1.2 eq.) were added. While stirring at 0°C, a mixture containing 0.01 eq of each acyl chloride was added dropwise to the mixture. The mixture was stirred for 2-3 h and allowed to gradually warm to 23°C. The mixture was then concentrated *in vacuo* and an LC-MS/MS sample was prepared.

**Esterification (c).** A carboxyl compound (1 eq.) was dissolved in ethanol in a round bottom flask with a magnetic stir bar. To the solution, a few drops of sulfuric acid were added. The solution was refluxed in an oil bath for 2-3 h and allowed to gradually warm to 23°C. The mixture was then concentrated *in vacuo* and an LC-MS/MS sample was prepared.

**Esterification (d).** A carboxyl compound (1 eq.) was dissolved in 2.5 mL of DMF in a scintillation vial with a magnetic stir bar. To the solution,10 eq. of Ghosez's Reagent was added and stirred until the reaction mixture turned yellow. Then 2 eq. of neat pyridine, and the OH containing compounds was added. The reaction mixture was stirred for 2-3 hr. The mixture was then concentrated *in vacuo* and an LC-MS/MS sample was prepared.

**Esterification (e).** A carboxylic acid compound (1 eq.) and 2.5 mL of anhydrous THF were added to a 20 mL scintillation vial with a magnetic stir bar. The solution was cooled to 0°C with an ice bath. To this solution, ethyl chloroformate (1.2 eq.) and neat $Et_3N$ (1.5ED eq.) were added. The solution was then stirred at 0°C for 1.5 hours. After 1.5 hours, a solution containing 0.01 eq of each alcohol was added dropwise to the activated carboxyl mixture. The mixture was then stirred for an additional 2-3 h at 23°C. The mixture was then concentrated *in vacuo* and an LC-MS/MS sample was prepared.

**Acetylation (a).** In a 20 mL scintillation vial, 1 eq. of the compound was added along with 2.5 mL of anhydrous DCM and a magnetic stir bar. The solution was cooled to 0°C in an ice bath, and 1.2 eq of anhydrous sodium carbonate was added along with 0.3 eq. of DMAP. Then, 1.2 eq. of acetic anhydride was added to the reaction mixture dropwise. The mixture was then stirred for another hour at 0°C, then allowed to warm to 23°C and stirred overnight. The mixture was then concentrated *in vacuo* and an LC-MS/MS sample was prepared.

**Acetylation (b).** To a 20 mL scintillation vial, 1 eq. of an alcohol compound was added along with 2.5 mL of anhydrous DCM. and a magnetic stir bar. Then, 2 eq. of EDC, 2 eq. of acetic acid, and 3 eq. of triethylamine were added to the mixture. The reaction was then stirred at 23°C for 5-6 hours. The mixture was then concentrated *in vacuo* and an LC-MS/MS sample was prepared.

**Sulfation.** In a 20 mL scintillation vial, 1 eq. of an alcohol was added along with 2.5 mL of anhydrous DCM and a magnetic stir bar. Then, 1.5 eq. of pyridine sulfur trioxide was added under dry argon. The reaction was then stirred for 1-3 hours at 23°C The mixture was then concentrated *in vacuo* and an LC-MS/MS sample was prepared.

**Methylation.** In a 20 mL scintillation vial, 1 eq. of an alcohol, carboxylic acid, or amine was added along with 2.5 mL of anhydrous DMF. Then, 1.5-2 eq. of sodium carbonate and 1.5-2 eq. of methyl iodide were added. The reaction was then left to stir overnight at 23°C. The mixture was then concentrated *in vacuo* and an LC-MS/MS sample was prepared.

**Oxidation.** In a 20 mL scintillation vial, 1 eq. of an alcohol was added along with 2.5 mL of DCM. Then, 0.04 eq. of TEMPO, 0.1 eq. of NaBr, and 2 eq. of sodium hypochlorite were added dropwise over 15 minutes. The reaction was then stirred for 2-3 hours. The mixture was then concentrated *in vacuo* and an LC-MS/MS sample was prepared.

**Oxidation of PUFAs** using $H_2O_2$. 1.4 eq. of free fatty acids were added to a 20 mL scintillation vial with 1mL of ethanol. The solution was then mixed with 1 eq. of 120 mM hydrogen peroxide in water solution. After 30 minutes, the reaction was divided into two 1 mL aliquots. In the first, 0.1 mL of 1M KOH in water was added. This aliquot was then incubated for 30 minutes at 40°C in a tube rotator. Afterwards, 100 µL of formic acid was added to obtain an acidic pH of 4. In the second aliquot, 1 mL of acidified water with formic acid (pH=4) was added.

**Boc deprotection (a).** A boc protected compound (1 eq.) was dissolved in 2 ml of Dioxane and cooled to 0°C with an ice bath.10 eq. of 4M HCl in Dioxane solution was added dropwise while stirring 0°C. The reaction was allowed to warm up to 23°C and stirred for 1-3 hours at room temperature. The reaction mixture was then concentrated *in vacuo* and an LC-MS/MS sample was prepared.

**Boc deprotection (b).** A boc protected compound (1 eq.) was dissolved in 2 ml of DCM and cooled to 0°C with an ice bath. Then, 10 eq. of TFA was added dropwise while stirring 0 °C. The reaction was then allowed to warm to 23°C and stirred for an additional 1-3 hours. The reaction mixture was then concentrated *in vacuo* and an LC-MS/MS sample was prepared.

**Glucuronidation**. Each substrate (30 mg, 1 eq.) was dissolved in toluene (3 mL). Acetobromo-α-D-glucuronic acid methyl ester (300 mg, 10 eq.) and silver carbonate (290 mg) were then added sequentially. The mixture was stirred vigorously and maintained under controlled heating for different time intervals 24 h or 3 days at 75°C, and up to 5 days at 90°C depending on the substrate, to achieve optimal conversion. After completion, the reaction mixture was cooled to 23°C, filtered to remove solids, and concentrated under reduced pressure. The resulting residue was redissolved in methanol (0.5 mL) and an equal volume of 1 M aqueous LiOH, and the solution was stirred at room temperature for 28 h. The solvent was then evaporated, and the mixture was neutralized with 1 M acetic acid (0.5 mL). The crude products were subjected directly to LC–MS/MS analysis.

**Reductive amination.** An amine compound (1 eq.) and a carbonyl compound (1 eq.) were mixed in anhydrous DCM (5 mL) and then treated with sodium triacetoxyborohydride (1.4 eq.) and AcOH (1 eq.). The mixture was stirred at 23 °C under argon for 24 h. The reaction mixture was then quenched by adding 1 M NaOH, and the product was extracted with DCM. The extract was then washed with brine and dried with $Na_2SO_4$. The solvent was then evaporated.

**Hydrolysis**. A mixture of methyl esters (1 eq.) was dissolved in THF:H2O (5 mL: 5 mL) and cooled to 0°C with an ice bath for 15 min before adding solid LiOH (5 equiv) in one portion. The reaction was then allowed to warm to 23°C and stirred 12 h. Excess hydroxide was then quenched with 1 M HCl. The product was extracted with ethyl acetate (5 mL × 3) and the combined organic extracts were washed with brine (5 mL), dried over sodium sulfate, filtered, and concentrated *in vacuo*.

**Ring opening.** In a 250 ml round bottom flask with an inert argon atmosphere, δ-decalactone (1 eq.) And sodium ethoxide (21% ethanol solution, 1.2 eq.) were added while stirring at room temperature.The reaction was left to stir overnight, then glacial acetic acid (1 eq.) was added to neutralize the NaOEt. The mixture was then concentrated under reduced pressure. Saturated brine (100 mL) was then added to the obtained residue, and the mixture was extracted with DCM (100 ml). The organic layer was then washed with saturated aqueous sodium bicarbonate solution (100 ml) and saturated brine (100 mL). The organic layer was then dried over anhydrous sodium sulfate and concentrated under reduced pressure.

**Acid chloride formation.** To a solution containing a carboxylic acid (1 eq.) in anhydrous Dichloromethane (DCM) (2.5 ml) oxalyl chloride was added(2 eq), followed by 2 drops of DMF. The mixture was stirred at 23°C for 30 min, then excess oxalyl chloride and dichloromethane were removed *in vacuo.*

**Glycosylation**. To a solution of a hydroxyl (1 eq.) containing compound in 3 mL of toluene, and (10 eq.) of acetobromo-sugar methyl ester was added, followed by the addition of (2 eq.) of silver carbonate. The reaction mixture was refluxed and stirred at 75°C for more than 24 hours. After reflux, the mixture was cooled to room temperature, filtered, and evaporated to dryness under vacuum. The resulting solid residue was dissolved in 10 mL of methanol, and (6 eq.) of 1 M aqueous LiOH were added. The mixture was stirred at 23°C for 28 hours. After the solvent removal, the product was neutralized with 0.5 mL of 1 M acetic acid. The reaction crudes were injected into the LC-MS/MS system and used for further analysis.
